# Lesion-remote astrocytes govern microglia-mediated white matter repair

**DOI:** 10.1101/2024.03.15.585251

**Authors:** Sarah McCallum, Keshav B. Suresh, Timothy Islam, Ann W. Saustad, Oksana Shelest, Aditya Patil, David Lee, Brandon Kwon, Inga Yenokian, Riki Kawaguchi, Connor H. Beveridge, Palak Manchandra, Caitlin E. Randolph, Gordon P. Meares, Ranjan Dutta, Jasmine Plummer, Simon R.V. Knott, Gaurav Chopra, Joshua E Burda

## Abstract

Spared regions of the damaged central nervous system undergo dynamic remodeling and exhibit a remarkable potential for therapeutic exploitation. Here, lesion-remote astrocytes (LRAs), which interact with viable neurons, glia and neural circuitry, undergo reactive transformations whose molecular and functional properties are poorly understood. Using multiple transcriptional profiling methods, we interrogated LRAs from spared regions of mouse spinal cord following traumatic spinal cord injury (SCI). We show that LRAs acquire a spectrum of molecularly distinct, neuroanatomically restricted reactivity states that evolve after SCI. We identify transcriptionally unique reactive LRAs in degenerating white matter that direct the specification and function of local microglia that clear lipid-rich myelin debris to promote tissue repair. Fueling this LRA functional adaptation is *Ccn1*, which encodes for a secreted matricellular protein. Loss of astrocyte CCN1 leads to excessive, aberrant activation of local microglia with (i) abnormal molecular specification, (ii) dysfunctional myelin debris processing, and (iii) impaired lipid metabolism, culminating in blunted debris clearance and attenuated neurological recovery from SCI. *Ccn1*-expressing white matter astrocytes are specifically induced by local myelin damage and generated in diverse demyelinating disorders in mouse and human, pointing to their fundamental, evolutionarily conserved role in white matter repair. Our findings show that LRAs assume regionally divergent reactivity states with functional adaptations that are induced by local context-specific triggers and influence disorder outcome.

Astrocytes tile the central nervous system (CNS) where they serve vital roles that uphold healthy nervous system function, including regulation of synapse development, buffering of neurotransmitters and ions, and provision of metabolic substrates^1^. In response to diverse CNS insults, astrocytes exhibit disorder-context specific transformations that are collectively referred to as reactivity^2–5^. The characteristics of regionally and molecularly distinct reactivity states are incompletely understood. The mechanisms through which distinct reactivity states arise, how they evolve or resolve over time, and their consequences for local cell function and CNS disorder progression remain enigmatic.

Immediately adjacent to CNS lesions, border-forming astrocytes (BFAs) undergo transcriptional reprogramming and proliferation to form a neuroprotective barrier that restricts inflammation and supports axon regeneration^6–9^. Beyond the lesion, spared but dynamic regions of the injured CNS exhibit varying degrees of synaptic circuit remodeling and progressive cellular responses to secondary damage that have profound consequences for neural repair and recovery^10,11^. Throughout these cytoarchitecturally intact, but injury-reactive regions, lesion-remote astrocytes (LRAs) intermingle with neurons and glia, undergo little to no proliferation, and exhibit varying degrees of cellular hypertrophy^7,12,13^. The molecular and functional properties of LRAs remain grossly undefined. Therapeutically harnessing spared regions of the injured CNS will require a clearer understanding of the accompanying cellular and molecular landscape.

Here, we leveraged integrative transcriptional profiling methodologies to identify multiple spatiotemporally resolved, molecularly distinct states of LRA reactivity within the injured spinal cord. Computational modeling of LRA-mediated heterotypic cell interactions, astrocyte-specific conditional gene deletion, and multiple mouse models of acute and chronic CNS white matter degeneration were used to interrogate a newly identified white matter degeneration-reactive astrocyte subtype. We define how this reactivity state is induced and its role in governing the molecular and functional specification of local microglia that clear myelin debris from the degenerating white matter to promote repair.

## Results

### Molecular dissection of LRAs after SCI

LRAs exhibit varying degrees of hypertrophy and intermingle with viable neurons, glia and neural circuitry throughout cytoarchitecturally intact regions of the injured CNS^7^ (Fig. 1a). Whether LRA reactivity evolves or resolves over time and how this form of reactivity differs from BFAs is unclear. We addressed these questions first by broadly examining injury-reactive gene expression dynamics of LRAs in a mouse model of anatomically and functionally incomplete SCI (iSCI). After iSCI, spared regions of the injured spinal cord rostral and caudal to the lesion undergo synaptic circuit reorganization that re-establish brain-cord communication and gives rise to recovery of locomotor behavior^14,15^ (Extended Data Fig. 1a). Concurrently, discrete spared white matter regions undergo widespread secondary axon degeneration, which gives rise to chronic gliosis and inflammation^16–18^. We performed bulk RNA sequencing of astrocyte-specific ribosome-associated mRNA (RiboTag^19^) and whole tissue mRNA from spared tissue regions rostral and caudal to the lesion epicenter at multiple post-injury timepoints. These time points reflect distinct phases of functionally meaningful neuroplasticity and locomotor recovery after iSCI^15^(Fig. 1b; Extended Data Fig. 1a). Thus, we could interrogate LRA transcriptional dynamics associated with post-traumatic neuroplasticity, inflammation and neurological recovery.

**Fig. 1.**
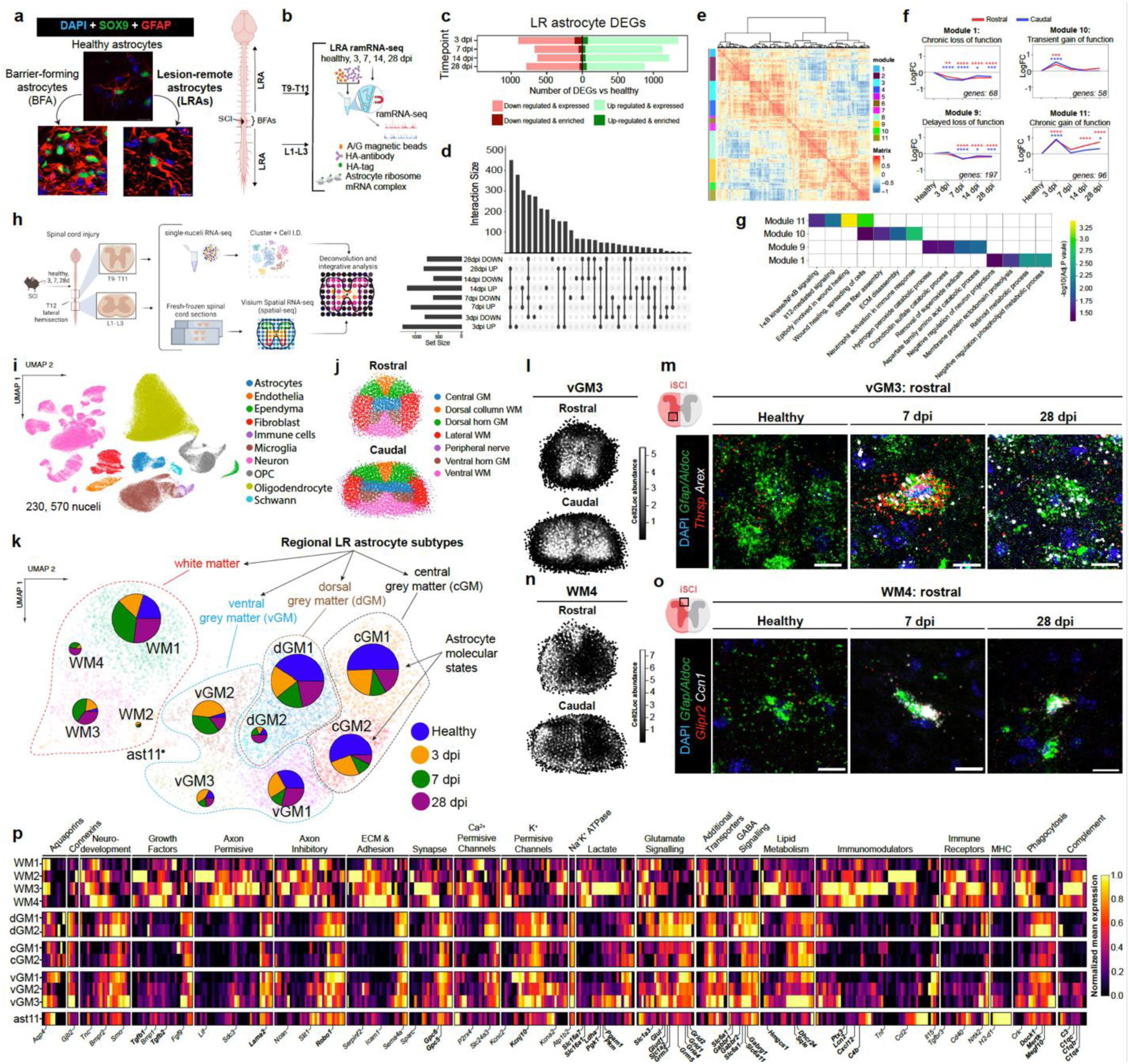
Spatiotemporal molecular decoding of SCI LRAs. **a**, Comparison of SCI BFAs and hypertrophic reactive LRAs after iSCI. **b**, SCI LRA RiboTag RNA-Seq schematic. **c**, **d,** Bar graph and upset plot illustrating LRA RiboTag RNA-Seq DEGs over time after iSCI, vs Healthy (FDR *P* ≤ 0.01) (n=4-6 mice/genotype/group). Rostral and caudal data are combined. **e**, Identification of temporally co-regulated LRA-enriched gene modules by Spearman correlation analysis. **f**, **g**, Line graphs illustrating temporally regulated gene expression of example LRA co-regulated gene modules with heat map of respective astrocyte relevant and enriched gene ontology terms. Temporal regulation patterns for rostral (red) and caudal (blue) are illustrated independently. (*P* ≤ 0.05, \*\**P* ≤ 0.002, \*\*\**P* ≤ 0.0002, \*\*\*\**P* ≤ 0.0001Two-way ANOVA with Tukeys). **h**, Schematic of combined snRNA-Seq and spatial transcriptomics approach for interrogating SCI reactive LRAs and neighboring cell types. **i**, UMAP of spinal cord cell types identified by snRNA-Seq for healthy and all post-injury time points, rostral and caudal (n=4-6 mice/timepoint/region). **j**, Diagram of intraspinal regions used in deconvolution of spatial transcriptomic data (n=4 mice/timepoint/region). **k**, UMAP constellation plot of healthy and iSCI astrocytes illustrating neuroanatomically restricted LRA subtypes and their distinct molecular states. **l**, Spatial transcriptomic characterization of vGM3 astrocytes illustrates restricted positioning in ventral horn grey matter. **m**, High magnification of vGM3 markers *Arex* and *Thrsp* in *Gfap^+^*/*Aldoc^+^* ventral horn grey matter *astrocytes*. **n**, Spatial transcriptomic characterization of WM4 astrocytes illustrates restricted unilateral white matter expression. **o**, High magnification of vGM3 markers shows unilateral expression of *Glipr2* and *Ccn1* in *Gfap^+^*/*Aldoc^+^ white matter astrocytes*. **p**, Heat map showing relative expression of functionally related genes across regionally restricted LRA molecular states. Scale bar, 10 µm.

Analysis of differentially expressed genes (DEGs) uncovered large and persistent alterations in astrocyte molecular profile that progress over time following injury, many of which were significantly astrocyte-enriched (Fig. 1c-g; Supplementary Information 1). Remarkably, SCI LRAs regulate only 15.4% of DEGs previously identified for SCI BFAs ^8^ (Extended Data Fig. 1b). LRAs from rostral and caudal regions exhibited only minimal transcriptomic differences (Extended Data Fig. 1c). To define temporally regulated genetic programs of LRA reactivity, we performed a gene-gene correlation analysis of LRA-enriched DEGs (Fig. 1e, f). This analysis identified eleven modules of co-regulated genes that exhibit divergent patterns of temporal regulation (i.e. transient, delayed or chronically altered expression) and distinct functional pathway enrichment (Fig. 1e-g; Extended Data Fig. 1e).

Next, we examined whether the transcription factor Signal transducer and activator of transcription 3 (STAT3), a well-established master regulator of hypertrophic astrocyte reactivity ^5,12,20^, is necessary for reactive alterations in LRA molecular profile. RiboTag RNA-Seq of iSCI LRAs from mice with inducible, astrocyte-specific *Stat3* deletion (*Stat3*cKO) showed that loss of STAT3 dramatically attenuates injury-reactive LRA gene expression, with ∼50% of WT LRA DEGs no longer modulated in astrocytes from *Stat3*cKO iSCI mice (Extended Data Fig. 1d; Supplementary Information 2). These results are in line with previous reports linking STAT3 to reactive transcriptional alterations in BFAs^5,8^. Together, these findings show that: (i) LRAs within spared regions of the injured spinal cord undergo a profound degree of transcriptional reprogramming that evolves over time after iSCI; and the majority of injury-reactive alterations in LRA gene expression are (ii) largely distinct from SCI BFAs and (iii) dependent on phosphorylation-dependent STAT3 signaling.

### SCI LRAs exhibit spatiotemporal molecular heterogeneity

In contrast to BFAs, LRAs tile anatomically and functionally discrete white and grey matter domains of grossly intact spinal cord regions^7^. Previous work has uncovered astrocyte regional heterogeneity within the healthy spinal cord^21–23^. Whether these regionally distinct astrocyte subtypes acquire unique reactivity states has not been investigated. We reasoned that since transcriptional profiles of reactive astrocytes are highly context-dependent^2–5^, LRAs from different neuroanatomical microenvironments of the injured cord may manifest divergent reactivity states. To test this hypothesis, we performed single-nuclei RNA-Seq (snRNA-Seq) and spatial transcriptomics on equivalent spared regions of the injured adult mouse spinal cord rostral and caudal to an iSCI lesion at 3, 7 and 28 days post-injury (dpi) (Fig. 1h-j). snRNA-Seq captured gene expression profiles of 230,570 cells, detecting all major CNS cell types in the healthy and injured spinal cord, but with differences in molecularly-defined cell states at different times post-injury (Fig. 1i; Extended Data. Fig. 2a-d).

We identified 12 distinct astrocyte molecular states, the relative proportions of which varied from healthy to injured, and across post-injury time points (Fig. 1k). snRNA-seq profiles were used to deconvolve spatial transcriptomic data and map astrocyte molecular states to their native intraspinal anatomical location (Fig. 1h, j; Extended Data Fig. 2e-i)^24^. This approach revealed that distinct snRNA-Seq astrocyte subtypes mapped to discrete anatomical regions within the white matter and grey matter of the healthy and injured spinal cord (Fig. 1k-o). Multiple grey matter astrocyte subtypes with discrete spatiomolecular profiles along the dorsoventral axis of the spinal cord were also defined. (Fig. 1k-m; Extended Data Fig. 3a-h). This separation of spinal cord astrocytes in functional neuroanatomical space paralleled marked transcriptional divergence, suggesting that region-specific injury-reactive alterations in astrocyte molecular state may have circuit-specific consequences (Fig. 1p; Supplementary Information 3).

### Decoding LRA reactivity states after SCI

We discovered multiple hypertrophic LRA reactivity states with characteristically elevated expression of intermediate filament genes (*Gfap*, *Vim)* and that exhibit unique spatiotemporal profiles (Extended Data Fig. 3a, i). LRAs in the sensory laminae of the dorsal horn grey matter (dGM1/2) demonstrated a notable shift in molecular profile towards a dGM2 molecular state at 28 dpi (Fig. 1k; Extended Data Fig. 3b-f). Relative to astrocytes in the healthy dorsal horn grey matter (dGM1), the chronic dGM2 astrocyte reactivity state is characterized by elevated expression for glutamate Transporter-1 (*Glt1*), developmentally regulated metabotropic glutamate receptor 5 (*Grm5*), and multiple ionotropic AMPA glutamate receptors (*Gria4, Grid1, Grid2*). dGM2 astrocytes also upregulate synaptogenic glypicans^25,26^ (*Gpc5*, *Gpc6*) and genes implicated in phagocytic clearance of synapses and cellular debris (*Megf10, Mertk, Dock1*) (Fig. 1p; Supplementary Information 3). LRAs that intermingle with motor neuron pools and associated interneurons within the ventral grey matter (vGM1/2/3) displayed a robust, but transient increase in vGM2 and vGM3 molecular state representation during acute and subacute post-injury timepoints (Fig.1k-m; Extended Data Fig. 3g). Relative to astrocytes in the healthy ventral grey matter (vGM1), vGM2/3 astrocytes dynamically upregulate metabotropic glutamate receptors *Grm3* and *Grm5*. C1q (*C1qa/b/c*) and *C4b* are also upregulated, implicating these astrocytes in complement-mediated synaptic circuit remodeling of spinal locomotor centers after SCI^27^. Intra-regional reactive astrocyte heterogeneity is underscored by vGM2 vs vGM3 transcriptional differences. For example, whereas LRAs acquiring a vGM2 reactivity state *downregulate* GABA transporters (*Gat1, Gat3*) and the primary astrocyte inward rectifying K^+^ channel Kir4.1 (*Kcnj10*), vGM3 LRAs *upregulate* expression of these genes. vGM3 astrocytes also exhibited distinctly elevated expression of GABA receptor subunits (*Gabbr1, Gabbr2, Gabrg1*), Glutamate uptake and metabolism genes (*Slc1a2, Slc1a3, Glul, Glud1*) and key sterol metabolism genes (*Hmgcs1, Dhcr24, Sqle*)(Fig.1p; Supplementary Information 3). Thus, LRAs in the grey matter acquire region-specific hypertrophic reactivity states (dGM2, vGM2, vGM3) with likely circuit-specific and functional consequences.

Hypertrophic white matter astrocytes exhibiting WM2/3/4 molecular states are restricted to the injured spinal cord and exhibit a lesion ipsilateral regional identity(Fig. 1k, n, o; Extended Data Fig. 2e, and 3a, h). White matter LRA transcriptomic profile evolves over time, with a greater proportion WM2/3 LRAs at acute and subacute time points, transitioning to a WM4 molecular state in the chronic post-injury phase (Fig.1k). Relative to healthy white matter astrocytes (WM1), WM2/3/4 LRAs displayed indicators of metabolic plasticity, namely widespread upregulation of lactate metabolism/transport (e.g. *Mct1, Mct2, Ldha*) and glycolysis genes (e.g. *Pgam1, Pgk1, Pkm*), which may underly astrocyte-mediated alterations in axonal energy metabolism in lesion-remote white matter (Fig. 1p; Supplementary Information 3). Remarkably, we determined that WM2/3/4 LRAs showed persistently elevated expression of immune and inflammamodulatory genes, potentially implicating them in chronic white matter inflammation and repair (e.g. *Lcn2, Cxcl12, Ptx3*, *Tgf-β1, Tgf-β2)*.

### Injury-reactive WM3/4 LRAs neighbor debris clearing microglia

Transition to a reactivity state may confer a gain-of-function that alters astrocyte-mediated cellular interactions, with adaptive or maladaptive consequences that modify disorder outcome^2–5,28^. Toward determining the potential functional implications of discrete LRA reactivity states, we moved next to define putative heterotypic cellular interactions involving region-specific LRAs and neighboring cell types. Non-negative matrix factorization (NMF) was applied on our deconvolved spatial transcriptomics data to identify co-occurring cell states across space and time (Fig.2a-c; Extended Data Fig. 4a). We identified 8 spatiotemporally distinct NMFs that predict diverse cellular landscapes containing distinct subtypes of LRAs and neighboring neurons, glia and endothelia, suggesting dynamic heterotypic cellular interactions (Fig. 2a-c; Extended Data Fig. 4). Evaluation of NMF 4 identified a collection of unilateral, white matter restricted cell populations only present in the injured spinal cord (Fig. 2a, b). This factor consisted of reactive white matter LRAs (WM3, WM4) and two white matter-restricted microglia subtypes (Mg2, Mg5)(Fig. 2c, d). In addition to their shared spatial pattern, Mg2 and Mg5 microglia also exhibited largely overlapping transcriptomic identities and were combined for subsequent analyses. (Fig. 2e-g; Extended Data Fig. 5). Relative to healthy microglia, Mg2/Mg5 microglia significantly upregulated genes involved in phagocytosis (*Lgals3*, *Cd68*, *Axl*), lipid metabolism (*Abca1*, *Abcg1*, *Plin2*, *Apoe*) and inflammation regulation (*Gpnmb, Igf1, Spp1*), and down-regulation of homeostatic microglia genes (*Tmem119*, *P2ry12*) (Fig. 2e, f; Extended Data Fig. 5d, e). There was also resemblance to multiple analogous types of phagocytic *‘white matter-associated microglia’ (WAM)* that localize to aging, damaged, and degenerating CNS white matter, where they have been implicated in dampening inflammation, cellular debris clearance and promoting tissue repair^29–33^(Fig. 2g). Similar microglia also populate white matter regions undergoing remodeling during development ^34–36^. Overlap in molecular signature with neurodegenerative disease-associated microglia (DAM)^37^, lipid-phagocytosing ‘microglia inflamed in MS’ (MIMS)^29^ and ‘trauma associated microglia’^38^ was also apparent (Fig. 2g). Thus, Mg2/5 microglial may represent a white matter inflammation and repair-associated state that is conserved across divergent CNS insults and disorders.

**Fig. 2.**
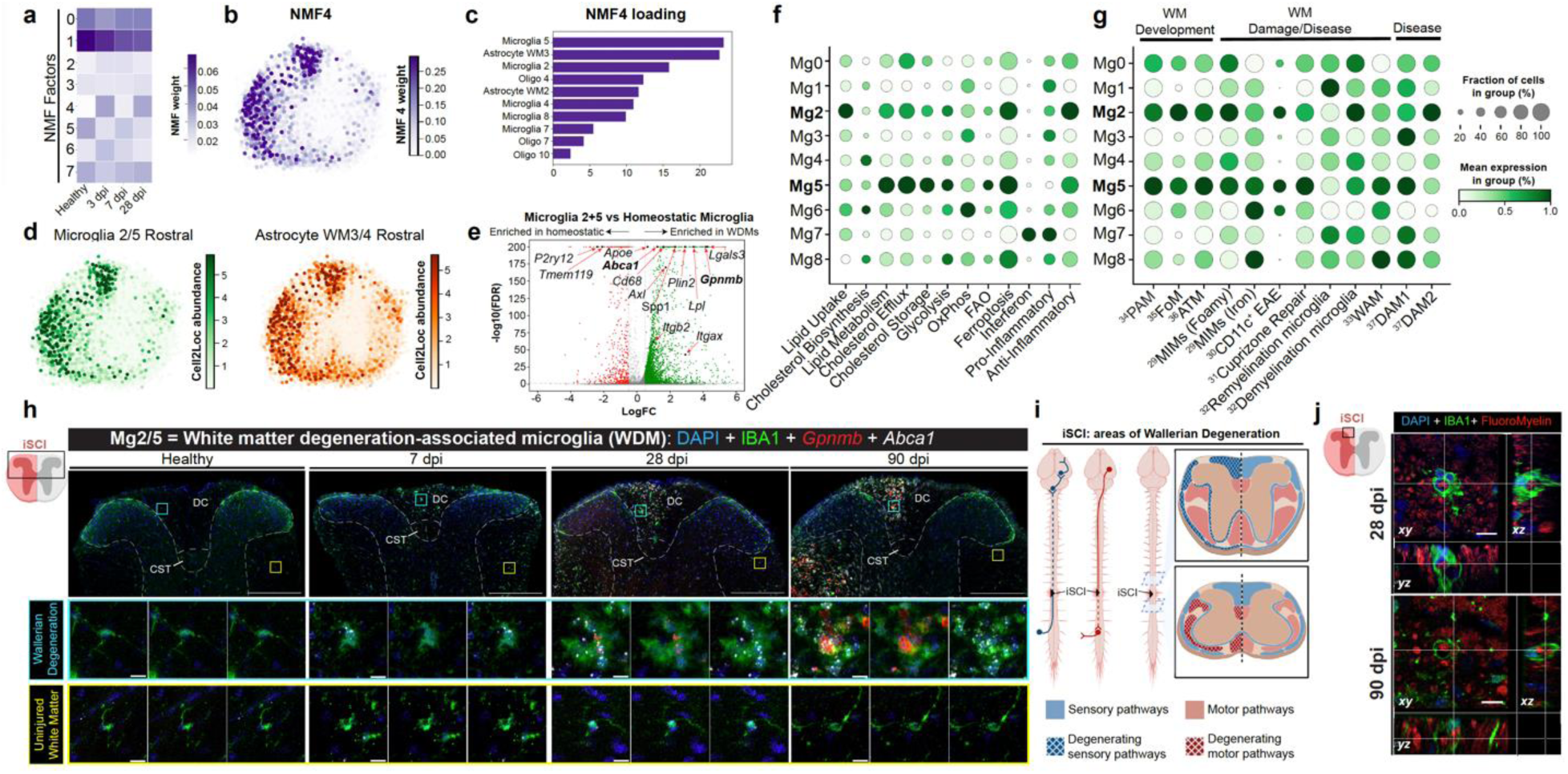
WM3/4 LRAs neighbor debris-clearing WDM in degenerating white matter. **a,** Temporally regulated NMF factors predicting injury-reactive transcriptional alterations in regionally co-occurring cell states. b, c, Spatial and cell identity loading profiles of NMF 4 reveals unilateral enrichment for WM3/4 LRA and Mg2/Mg5 microglia. d, Spatial transcriptomic characterization of Mg2 and Mg5 microglia illustrating unilateral white matter enrichment. e, Volcano plot of DEGs in Mg2/5 microglia vs healthy microglia (Mg1) (FDR P ≤ 0.01). f, g, Dot plot of mean normalized expression of metabolism- and previously published microglia state-associated molecular signatures in iSCI lesion-remote microglia clusters. h-j, Expression of Mg2/5 marker genes (*Gpnmb, Abca1*) designates ‘WDM’ microglia that localize to Wallerian degenerating white matter regions and assemble into multi-cellular nodules that phagocytose myelin debris. Scale bars: h, low magnification, 250 µm; j and h inset, 10 µm.

We determined that molecular markers of the Mg2/5 snRNA-Seq profile (*Gpnmb, Abca1*) specifically correspond to white matter microglia that gradually assemble into multi-cellular nodules within the lesion ipsilateral white matter and persist for at least three months after iSCI (Fig. 2h). Notably, appreciable numbers of WDM multi-cellular nodules are not apparent until after 7 dpi and increase in density thereafter (Fig. 2h). After SCI, axons in lesion-remote white matter undergo Wallerian degeneration, resulting in the generation of myelin and axon debris, and the recruitment of numerous debris-clearing microglia that arrange as multi-cellular ‘nodules’^17,39^. Correspondingly, we find that Mg2/5 microglia rostral to the iSCI lesion are restricted to Wallerian degenerating sensory tracts of the dorsal column white matter (DC), but mostly absent from the descending motor fibers of the corticospinal tract (CST), which are severed after iSCI, but do not undergo Wallerian degeneration in this region (Fig. 2h, i). We find also that Mg2/5 microglia nodules phagocytose myelin debris in Wallerian degenerating axon tracts (Fig. 2j). Therefore, we refer to these cells collectively as *white matter degeneration-associated microglia* (WDM).

Collectively, these results demonstrate that WDM: i) are found within Wallerian degenerating white matter regions of the injured spinal cord, associate into multi-cellular nodules and phagocytose myelin debris, ii) exhibit a distinct molecular signature typified by the upregulation of phagocytosis, lipid metabolisms and inflammomodulatory genes, and iii) may coexist with transcriptionally distinct reactive white matter astrocyte subtypes (WM3/4).

### WDMs intermingle with *Ccn1^+^* LRAs

Although some foundational properties of debris-clearing microglia are characterized in white matter damage and disease^31,33,40–46^, the cellular interactions shaping microglia responses, and how these impact debris clearance efficacy, inflammation regulation, repair or recovery after CNS injury are not well defined. Results from our NMF analysis suggested a remarkable regional relationship between distinct hypertrophic white matter astrocytes (WM3/4) and WDMs (Fig. 2b, c). Accordingly, we carried out NicheNet analysis ^47^ to identify ligand-receptor mediated pathways of communication between hypertrophic white matter astrocytes and WDM (Fig. 3a). Multiple regulators of extracellular proteolysis (A 2M, ADAM12, ADAM15, PLAT), matricellular proteins (TNC, CCN1) and inflammomodulatory molecules (C3, CXCL12, LGALS3) were among the temporally conserved astrocyte-derived ligands most strongly upregulated after iSCI.

**Fig. 3.**
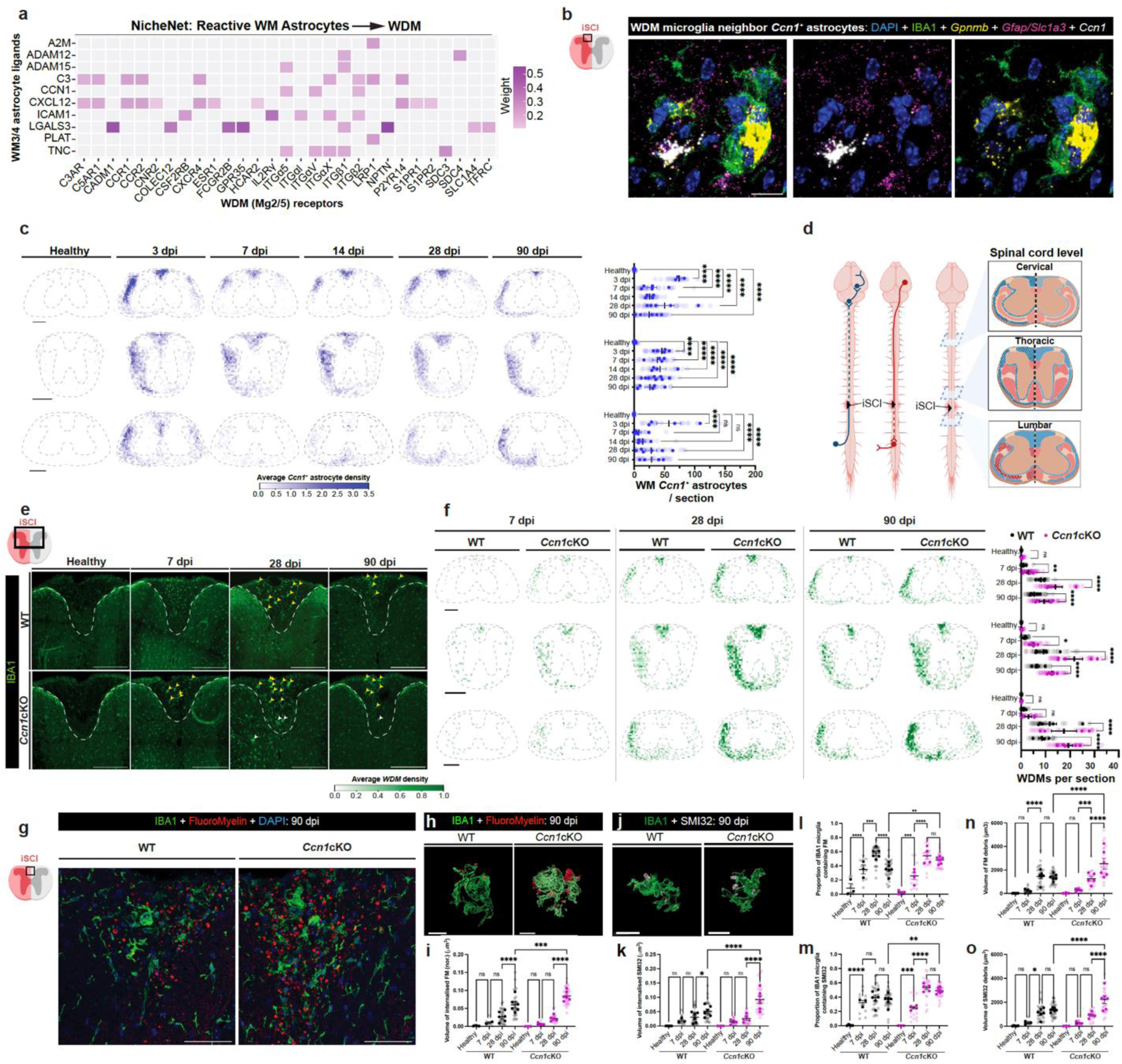
White matter degeneration-reactive astrocytes secrete CCN1 to regulate WDM nodule formation and cell debris clearance. **a**, Putative ligands from WM3/4 LRAs (senders) and predicted receptors enriched in WDM (Mg2/5) (receivers) (NicheNet). **b**, High magnification of predicted WM3/4 ligand *Ccn1*, shows selective expression in LRAs that neighbor WDM nodules in degenerating white matter. **c**, Aligned average density plots of *Ccn1*-expressing astrocytes illustrating regional and intraspinal relationship between *Ccn1*^+^ LRAs and anatomically defined zones of Wallerian degeneration. Plot shows quantification of *Ccn1^+^* astrocytes per timepoint per region. n=4-7 mice/spinal cord region; 181-204 sections per spinal cord region; 85-110 sections/time point; 580 sections total. **d**, Schematic of iSCI lesion-remote spinal cord regions evaluated in spatiotemporal analysis of *Ccn1*-expressing astrocytes. **e**, WDM nodules in Wallerian degenerating dorsal white matter (mid-thoracic cord) from WT and *Ccn1*cKO cord, over time, after iSCI. Yellow arrowheads represent WDMs in Wallerian degenerating regions, white arrowheads represent WDMs in aberrant areas. **f**, Aligned average density plots of WDM nodules illustrating accelerated, excessive and spatially aberrant nodule formation in *Ccn1*cKO spinal cord. WDM Density plots further illustrate a Wallerian degeneration-associated regional pattern. Plot shows quantification of WDM nodules per timepoint per region. n=4-6 mice/spinal cord region/genotype; 113-146 sections/spinal cord region/genotype; 83-110 sections/time point/genotype; 366 sections total. **g**, Low magnification of FluoroMyelin^+^ myelin debris and IBA1^+^ WDM nodules in Wallerian degenerating dorsal white matter of iSCI lesion-remote spinal cord (mid-thoracic cord) in WT vs. *Ccn1*cKO. **h-k** High magnification 3D image showing IBA1^+^ WDM

Expression validation of top ranked putative white matter astrocyte-derived ligands determined that cellular communication network factor 1 (*Ccn1)* is prominently and specifically expressed by astrocytes within degenerating spinal cord white matter that intimately associate with nodules of myelin debris-clearing WDM (Fig. 3b; Extended Data Fig. 6a). *Ccn1* encodes for a multifunctional, secreted matricellular protein implicated in peripheral inflammation and wound healing^48,49^. CCN1 acts on target cells, including macrophages primarily through different integrin heterodimers, in a context-dependent manner^48–50^. Congruently, astrocyte CCN1 is predicted to target multiple WDM-enriched integrins, including ITGα5, ITGαv, and ITGβ2 (Fig. 3a). Indeed, we found that WDM upregulate ITGβ2 for at least 3 months after iSCI (Extended Data Fig. 6b).

*Ccn1*-expressing astrocytes are exceptionally rare in healthy spinal cord but become detectable as early as 3 days post-iSCI and persist for at least 90 dpi (Fig. 3c; Extended Data Fig. 6c). Notably, *Ccn1^+^* LRAs contained significantly elevated nuclear levels of transcriptional coactivator YAP1, a canonical indicator of active *Ccn1* transcription^51^ (Extended Data Fig. 6d). A spinal cord-wide, spatiotemporally resolved analysis of *Ccn1*-expressing astrocytes revealed an unequivocal intraspinal regional relationship between *Ccn1*^+^ astrocytes and anatomically defined zones of Wallerian degeneration (Fig. 3c, d). For example, *Ccn1*^+^ astrocytes rostral to the lesion are mainly restricted to Wallerian degenerating DC sensory tracts, while those in the dorsal white matter caudal to the lesion are within the Wallerian degenerating CST (Fig. 2h, i; Fig. 3c, d; Extended Data. Fig. 6e). Thus, the spatial pattern of *Ccn1*^+^ astrocytes is evidently consistent with WDM nodules (Fig. 2h). Notably, we found that in a crush SCI model, with *bilateral* WD, elicits *bilateral Ccn1* expression in white matter astrocytes (Extended Data. Fig. 6f).

*Together, these data demonstrate that, after SCI, a subset of hypertrophic reactive white matter LRAs i) rapidly and persistently express Ccn1, ii) are restricted to Wallerian degenerating white matter, and iii) neighbor myelin debris-clearing WDM nodules that express multiple predicted CCN1 integrin receptors*.

### Astrocyte CCN1 regulates WDM nodule formation and cell debris clearance

To elucidate the function of astrocyte-secreted CCN1, we examined WDM specification and function throughout lesion-remote Wallerian degenerating white matter after iSCI in young adult WT mice, and mice with conditional astrocyte-specific^52^ *Ccn1* gene deletion^53^ (*Ccn1*cKO) (Extended Data Fig. 7a-e). Given the sustained intimate spatial relationship between *Ccn1*^+^ astrocytes and WDM (Fig. 3b), we quantified nodule formation dynamics across the rostrocaudal expanse of the injured spinal cord (Fig. 3e, f). We determined that *Ccn1*cKO mice exhibit a significant 2-4-fold greater number of nodules than WT at all levels of the spinal cord, and at virtually all post-injury time points examined up to 90 dpi (Fig. 3e, f). Nodule formation was significantly accelerated in the *Ccn1*cKO white matter, as indicated by notable nodule formation as early as 7 dpi (Fig. 3e, f). Strikingly, rostral to the injury, WDM nodules were regularly present in the CST of *Ccn1cKO* animals despite infrequently occurring in the CST in WT animals (Fig. 3e, f; Extended Data Fig. 7f). *Ccn1cKO* iSCI spinal cords also contained greater numbers of nodules in the contralateral white matter, which normally undergoes relatively minimal degeneration in this model (Fig. 3f; Extended Data Fig. 7g). Curiously, grey matter nodules were also significantly more common in *Ccn1cKO* iSCI spinal cord, relative to WT (Extended Data Fig. 7h). Therefore, loss of astrocyte-secreted CCN1 results in accelerated, excessive and spatially aberrant activation of phagocytic white matter microglia.

Given the importance of WDM nodules in myelin debris clearance, we investigated whether loss of astrocyte CCN1 impacts the ability of WDM to phagocytose lipid-rich myelin and axon debris from the degenerating spinal cord white matter in WT and *Ccn1cKO* iSCI mice (Fig. 3g-o). By 90 dpi, *Ccn1*cKO microglia contained significantly greater volumes of internalized myelin and axon debris than their WT equivalents (Fig. 3h-k). Loss of astrocyte-secreted CCN1 resulted in a greater overall proportion of microglia containing myelin or axon debris at 90 dpi (Fig. 3l, m). Yet, provocatively, we observed that loss of astrocyte CCN1 also leads to significantly attenuated debris clearance, with chronic accumulation of myelin and axon debris (Fig. 3n, o). Together, these data suggest that astrocyte-secreted CCN1 regulates the nodule formation dynamics of WDM and is necessary for the clearance of myelin and axon debris from degenerating white matter.

### Astrocyte CCN1 regulates WDM molecular specification and lipid metabolism

Excessive intracellular accumulation of myelin debris-derived lipids and attenuation of myelin debris clearance has been linked to maladaptive alterations in microglia lipid metabolism^31,33,40,54^. Congruently, our results thus far demonstrate that loss of astrocyte CCN1 leads to amplified activation of debris-laden microglia nodules, but overall impaired debris clearance. Therefore, we moved next to determine whether astrocyte-secreted CCN1 regulates microglia lipid metabolism by performing an unbiased lipidomics analysis on whole-cell extracts of microglia from lesion-remote spinal cord regions undergoing Wallerian degeneration in WT and *Ccn1cKO* mice at 28 dpi (Fig. 4a; Extended Date Fig. 8). Though microglia lipidomes from the healthy WT or *Ccn1cKO* spinal cord were grossly similar (Fig. 4b; Extended Data Fig. 8e), we observed highly divergent injury-induced alterations in lipidomic profile across multiple lipid classes (Fig. 4c-e; Extended Data Fig.8b-g). For example, in contrast to WT, *Ccn1cKO* iSCI microglia show increased levels of Sphingomyelins (SM), which parallels lipid metabolic pathway predictions suggesting an increased conversion of ceramides to SM (Extended Data Fig. 8f, g). *Ccn1c*KO iSCI microglia contained significantly elevated levels of multiple lipid classes found in myelin and axonal debris, including phoshatidylethanolamine (PE), SM and phosphatidylcholine, echoing the elevated levels of internalized myelin debris observed *in vivo* (Fig. 4c-e; Extended Date Fig. 8g).

**Fig. 4.**
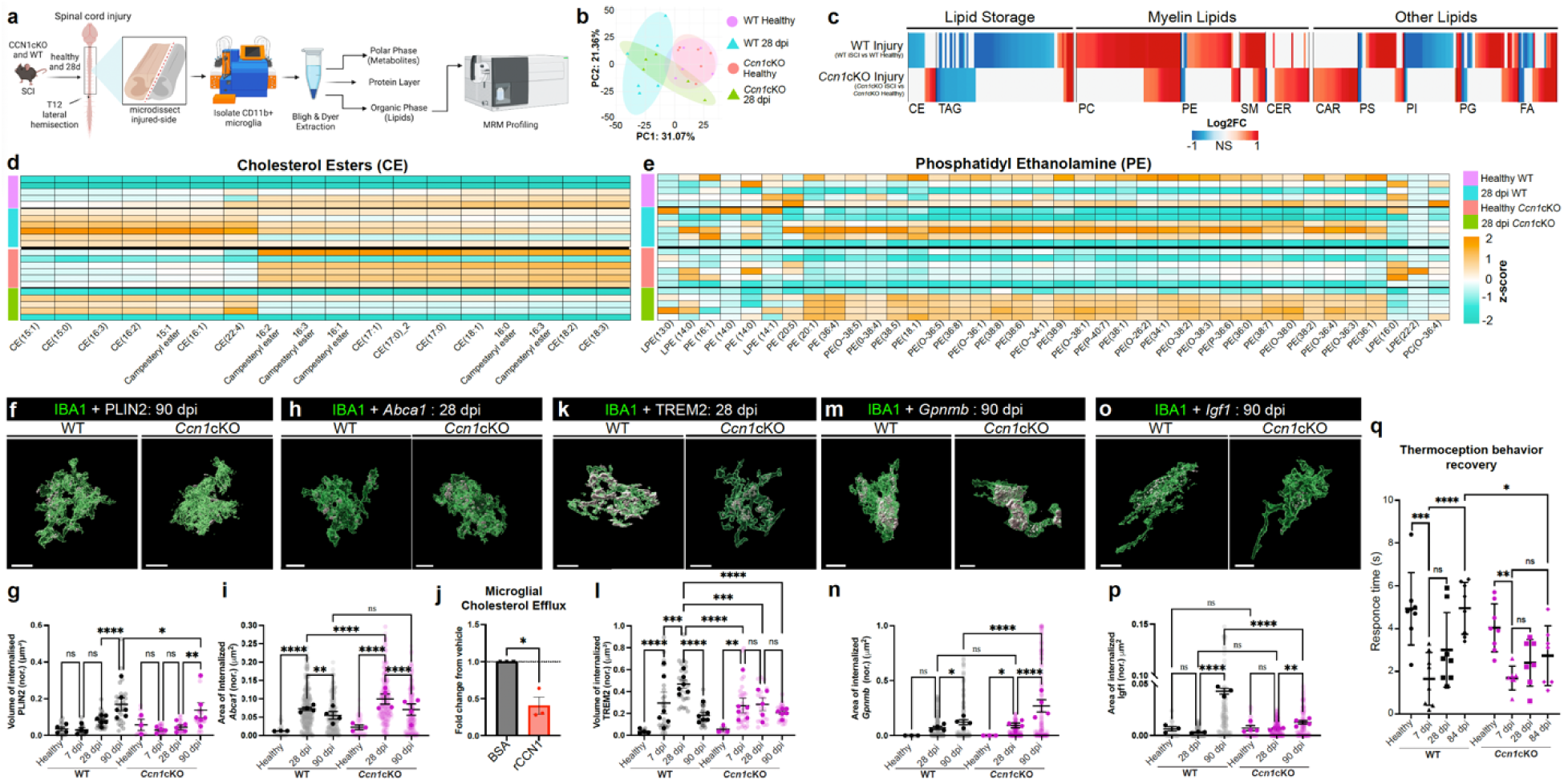
Astrocyte-secreted CCN1 regulates WDM molecular specification and myelin debris-associated lipid metabolism. **a**, iSCI lesion-remote microglia lipidomics schematic. A parasagittal spinal cord dissection enriched for microglia from Wallerian degenerating white matter. **b**, Comparison of lipidomic profiles of healthy and iSCI microglia from WT and *Ccn1*cKO in principle component space. iSCI microglia from WT and *Ccn1*cKO are clearly separable. **c**, Comparison of WT and *Ccn1*cKO significant injury-reactive alterations in microglia lipid profile, including lipid droplet- and myelin-associated lipid subtypes (Log_2_ fold-change, iSCI vs healthy, FDR *P* ≤ 0.01). Non-significantly altered lipid species show in white. **d**, **e**, Comparison of differentially expressed CE and PE lipid species in iSCI vs healthy microglia, from WT and *Ccn1*cKO spinal cords. **f-i**, High magnification 3D image and quantification of PLIN2^+^ lipid droplets and *Abca1* in WT and *Ccn1*cKO WDM from Wallerian degenerating white matter (PLIN2: n=3-6 mice/group; *Abca1*: n=3-4 mice/group). **j**, Cholesterol efflux measured from cultured primary mouse microglia following stimulation with CCN1 or vehicle (BSA) (n= 3 replicates from independent cultures conditions run in triplicate; Students t-test, \*\**P* ≤ 0.05)**. k-p**, High magnification 3D image and quantification of TREM2, *Gpnmb* and *Igf1* in WT and *Ccn1*cKO WDM from Wallerian degenerating white matter (TREM2: n=3-6 mice/group; *Gpnmb* and *Igf1* n=3-4 mice/group) **q**, Quantification of spontaneous recovery of left hind-paw cold thermoception over time after iSCI (n= 7-8 mice per group). CE: Cholesterol Esters, TAG: Triacylglycerol, PC: Phosphatidylcholine, PE: Phosphatidylethanolamines, EM: Sphingomyelin, CER: Ceramides, CAR: Carnitine, PS: Phosphatidylserine, PI: Phosphatidylinositol, PG: Phosphatidylglycerols, FA: Fatty Acids. Graphs show mean ± SEM. Opaque and transparent data points illustrate experimental replicate mean and counts from individual tissue sections/cells, respectively. Unless stated otherwise, \**P* ≤ 0.05, \*\**P* ≤ 0.002, \*\*\**P* ≤ 0.0002, \*\*\*\**P* ≤ 0.0001, Two-way ANOVA with Tukeys. Scale bars: 10μm.

Following phagocytosis, myelin-derived cholesterol is converted into cholesterol esters (CEs) and stored as lipid droplets, along with triacylglycerols (TAGs)^55^. Though our lipidomic analysis did not permit for the detection of free cholesterol, we found that *Ccn1cKO* iSCI microglia consisted of a significantly lower proportion of CEs and TAGs, relative to WT (Fig. 4c, d). Correspondingly, quantification of WDM intracellular perilipin 2 (PLIN2) revealed that *Ccn1*cKO WDM contain significantly fewer lipid droplets than their WT equivalents by 90 dpi (Fig. 4f, g; Extended Data Fig. 8h). Examination of WDM intracellular BODIPY^+^ inclusions, a common marker for lipid droplet associated neutral lipids^56^, further validated that *Ccn1*cKO WDM contain fewer lipid droplets than WT (Extended Data Fig. 8i-k).

In addition to being esterified and stored in lipid droplets, intracellular cholesterol exists also in a free state, which can be actively transferred back into the extracellular space via the ATP-binding cassette (ABC) transporters, ABCA1 and ABCG1. Excessive intracellular accumulation of cholesterol is known to drive upregulated expression of *Abca1* in macrophages and microglia^57,58^. We hypothesized that the accumulation of myelin debris-derived free cholesterol in *Ccn1*cKO microglia could result in amplified *Abca1* expression, which would favor cholesterol efflux over storage in lipid droplets. In alignment with previous work linking CCN1 to *Abca1* expression^59^, we observed that *Ccn1*cKO-derived WDM indeed express greater levels of *Abca1* than WT WDM at 28 dpi (Fig. 4h, i). Congruently, treatment of WT microglia with recombinant CCN1 significantly reduced cholesterol efflux *in vitro* (Fig. 4j). Thus, CCN1 is a bi-directional regulator of microglial cholesterol efflux.

Dysregulated lipid metabolism is associated with altered molecular specification of phagocytic microglia ^31,33,40,54^. Triggering receptor expressed on myeloid cells 2 (TREM2) is a cell surface immune receptor that binds numerous ligands, including myelin debris-derived lipids^40^. TREM2 signaling facilitates white matter microglia activation, drives myelin debris phagocytosis, and mediates transcriptional reprogramming necessary for metabolizing lipid-rich myelin debris^33,40^. Thus, we aimed next to determine whether differences in TREM2 levels may underlie the abnormal myelin debris management and lipid metabolism profile we observed in *Ccn1*cKO-derived WDM. In WT-derived WDM, we observed dynamic modulation of TREM2, which reached peak levels by 28 dpi and tapered significantly by 90 dpi. In contrast, TREM2 levels in *Ccn1*cKO-derived WDM plateaued by 7 dpi, remained unaltered to 90 dpi and failed to reach peak levels observed in WT cells (Fig. 4k, l; Extended Data Fig. 8l). Yet, WT and *Ccn1*cKO WDM showed equal levels of the lipid scavenger receptor lipoprotien lipase (LPL) (Extended Data Fig. 8m-o). We found also that *Ccn1*cKO-derived WDM exhibit significantly enhanced expression of *Gpnmb*, a potential indicator of lysosomal dysfunction^60^, and impaired expression of *Igf1*, both of which are central molecular indicators of the WDM state and encode for potent immunomodulatory proteins associated with debris clearance and white mater repair (Fig. 4m-p). Together, these results show that astrocyte-secreted CCN1 potently governs WDM debris processing via regulation of lipid metabolism, as well as associated molecular and functional specification.

### Astrocyte CCN1 promotes recovery of sensation after SCI

Failure to clear myelin debris can interfere with axon regeneration^61^ and restrict remyelination ^62,63^. Therefore, we examined whether impaired astrocyte *Ccn1* expression impacts on spontaneous recovery of sensorimotor function after iSCI. We initially employed a modified Basso mouse scale^64^ to compare recovery of locomotor function after iSCI in WT and *Ccn1*cKO mice, though no differences were observed (Extended Data Fig. 8p). iSCI severs the spinothalamic tract, which transmits information about pain, temperature and itch. We evaluated spinothalamic function to 84 days after iSCI by measuring hind-paw sensitivity to non-noxious cold stimuli^65^. In contrast to WT iSCI mice, who exhibited a full recovery of cold sensitivity, *Ccn1*cKO iSCI mice failed to recover cold thermoception (Fig. 4q). Therefore, astrocyte-secreted CCN1 not only regulates microglia-mediated white matter inflammation and repair, but is necessary for neurological recovery after SCI.

### Astrocyte *Ccn1* is induced by myelin degeneration

The nature of astrocyte-*extrinsic* mechanisms that trigger discrete reactivity states are poorly understood. Using *Ccn1* expression as a biomarker of a molecularly-distinct white matter astrocyte reactivity state, we moved next to determine the mechanism of its induction. Taken that astrocytes exhibiting a *Ccn1*-associated reactivity state localize to zones of white matter degeneration, we investigated whether myelin debris is sufficient to induce astrocytic *Ccn1* expression. We observed that intraspinal injection of CFSE-conjugated CNS myelin triggered robust astrocytic *Ccn1* expression (Fig.5a, b; Extended Data Fig. 9a). Notably, off-target injections placing myelin into the grey matter also generated *Ccn1*^+^ astrocytes, indicating that myelin damage, not the cellular regional microenvironment *per se*, mediates astrocyte *Ccn1* expression (Extended Data Fig. 9b).

Injections into white matter inherently damage axons and their associated myelin, obscuring whether astrocyte *Ccn1* expression is initiated by the degeneration of axons, myelin, or both. To tweeze apart this central mechanistic detail, we carried out intraplantar hind paw injection of saporin conjugated to either cholera toxin subunit B (CTB) or isolectin B4 (IB4), which selectively degenerate myelinated and non-myelinated primary sensory axons, respectively^66^(Fig. 5c). We then quantitatively assessed astrocyte *Ccn1* expression along degenerating central afferent fibers that innervate the spinal cord. Here, we observed that the degeneration of myelinated, but not non-myelinated axons resulted in significant astrocytic *Ccn1* expression (Fig. 5d-f). Moreover, the *Ccn1* expression induced by myelinated axon degeneration also preceded the arrival of multicellular nodules of IBA1^+^ microglia, which bear resemblance to myelin debris-clearing WDM of the injured spinal cord (Fig. 5e, g). We did not observe *Ccn1*^+^ astrocytes following mouse sciatic nerve crush, indicating that astrocyte *Ccn1* expression depends on myelin breakdown and not a generalized neuronal stress response (Extended Fig. 9c, d)

**Fig. 5.**
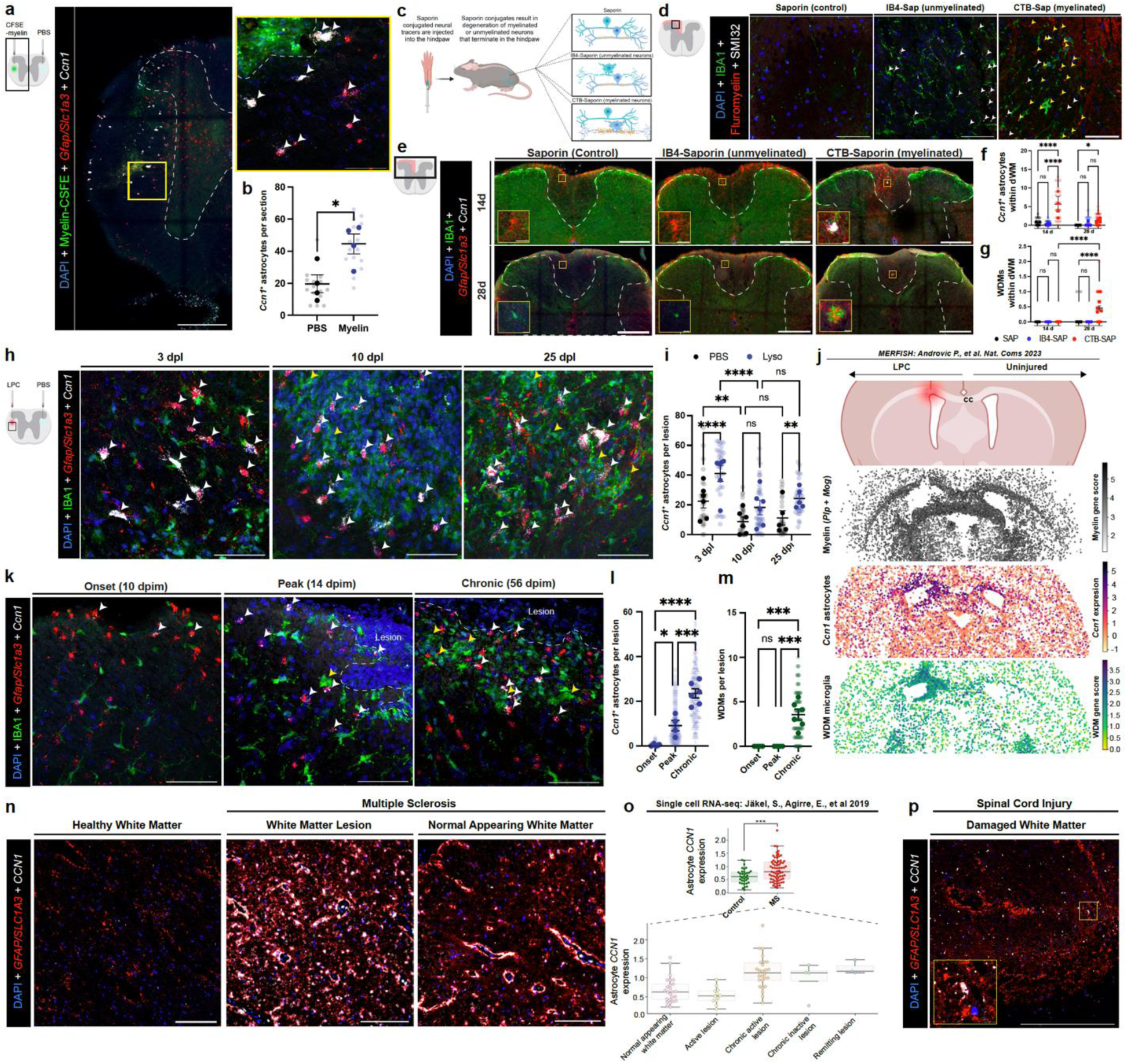
*Ccn1*^+^ astrocytes are induced by myelin damage and generated in diverse CNS white matter disorders in mouse and human. **a**, **b**, *Ccn1*^+^ astrocytes in mouse spinal cord lateral white matter following microinjection of CSFE-conjugated myelin and quantification relative to vehicle control (PBS)(n=4 mice/group; \**P* ≤ 0.05, Student’s t-test). **c**, Schematic of intraplanar saporin injection study for determining mechanism of astrocyte *Ccn1* induction. **d**, Fluoromyelin (yellow arrowheads) and SMI32 (white arrowheads) staining to detect myelin and axon degeneration following saporin injections. **e**-**g**, *Ccn1*^+^ astrocytes and IBA1^+^ microglia in spinal cord dorsal white matter following saporin-mediated neurodegeneration. *Ccn1^+^* astrocytes and WDM-like nodules are largely restricted to CTB-Saporin (n=3-4mice/group). **h**, **i**, *Ccn1*^+^ astrocytes (white arrowheads) and IBA1^+^ microglia/macrophages nodules (yellow arrowheads) in mouse spinal cord lateral white matter at 3, 10 and 25 days following microinjection of LPC. Quantification of *Ccn1*^+^ astrocytes relative to vehicle (PBS) (n=5-6 mice/group). **j**, spatial transcriptomic analysis correlating regions of LPC-demyelinated corpus callosum with *Ccn1*^+^ astrocytes and microglia exhibiting a WDM molecular profile^67^. **k**, *Ccn1*^+^ astrocytes (white arrowheads) intermingle with IBA1^+^ microglia/macrophages nodules (yellow arrowheads) neighboring mouse MOG_35-55_ EAE spinal cord white matter lesions. **l**, **m**, Quantification of *Ccn1*^+^ astrocytes and IBA1^+^ microglia nodules per EAE lesion (n=5-6 mice/group). **n**, *CCN1*^+^ astrocytes in a healthy individual, and MS white matter lesion and throughout normal appearing white matter in human spinal cord. **o**, Comparison of astrocyte *CCN1* expression in healthy and MS brain by single cell RNA-seq^69^. Top plot: control: 35 cells, MS samples: 70 cells; \*\*\**P* ≤ 0.0002, Studen’t t-test). **p**, *CCN1*^+^ astrocytes in lesion-remote Wallerian degenerating corticospinal tract white matter in human. Graphs show mean ± SEM. Opaque and transparent data points illustrate experimental replicate mean and counts from individual tissue sections, respectively. Unless stated otherwise, \**P* ≤ 0.05, \*\**P* ≤ 0.002, \*\*\**P* ≤ 0.0002, \*\*\*\**P* ≤ 0.0001, Two-way ANOVA with Tukeys. Scale bars: a, e, n, p: 250μm, a-inset,d,h,k: 50μm e-inset, p-inset: 10μm.

### *Ccn1^+^* astrocytes are a conserved feature of white matter damage in mouse and human

Results thus far suggested that astrocyte *Ccn1* expression may be a conserved response implicated in the regulation of phagocytic microglia in CNS white matter damage and degeneration. To explore this possibility, astrocyte *Ccn1* expression was examined in the context of demyelinating diseases and insults in mice and humans.

We quantified *Ccn1*-expressing astrocytes at 3, 10 and 25 days following focal demyelination by stereotaxic injection of lysolecithin into the mouse spinal cord white matter. *Ccn1*^+^ astrocytes were present by 3 days post-lesioning and persisted for at least 25 day thereafter, and neighbored microglia nodules (Fig. 5h, i; Extended Data Fig. 9 e-g). We also interrogated astrocyte *Ccn1* expression and molecular indicators of the WDM state from a recently published spatial transcriptomics dataset from lysolecithin demyelination in the mouse brain^67^. Here we determined, once again, that *Ccn1^+^* astrocytes and WDM microglia are intimately associated in demyelinated regions (Fig. 5j).

We next assessed astrocyte *Ccn1* expression in the spinal cords of mice with experimental autoimmune encephalomyelitis (EAE) at onset, peak and chronic disease (Extended Data 9h). Few *Ccn1*^+^ astrocytes were present at disease onset, localizing mainly to normal appearing ventrolateral white matter (Fig. 5k, l). In peak and chronic disease*, Ccn1^+^* astrocytes were prevalent, concentrated around inflammatory white matter lesions, and adjacent to microglia nodules (Fig.5k-m). Analysis of previously published RiboTag RNA-Seq data from mouse EAE spinal cord astrocytes^68^ further confirmed that demyelination-reactive astrocytes significantly upregulate *Ccn1* (Extended Data Fig. 9i).

Astrocyte *CCN1* expression was also evaluated in human archival spinal cord tissue from individuals with MS or SCI and neurologically healthy controls. *CCN1* expressing astrocytes were frequent in MS white matter, but rare in MS grey matter and healthy human spinal cord (Fig. 5n; Extended Data Fig. 9j, k). Interrogation of published single cell RNA-Seq from human MS and healthy brains provided further confirmation of our results^69^ (Fig. 5o). Corroborating our mouse iSCI results, *CCN1^+^* astrocytes were also observed throughout Wallerian degenerating white matter of the SCI-lesion remote human spinal cord, but not in neighboring grey matter (Fig. 5p; Extended Data Fig. 9l).

Together, these results demonstrate that astrocyte-secreted CCN1 expression is i) an evolutionarily conserved response of white matter damage-reactive astrocytes, ii) induced by myelin degeneration, and iii) implicated in the regulation of debris-clearing phagocytes across divergent forms of CNS white matter damage and disease in mice and humans.

## Discussion

The present work resolves multiple fundamental attributes of reactive LRAs, including i) how LRAs vary across specific neuroanatomical environments, ii) how the molecular phenotype of distinct subtypes evolves or resolves over time, iii) the roles these astrocytes play in local multicellular responses to CNS injury and repair, and iv) the cellular and molecular mechanisms that drive distinct states of LRA reactivity.

Astrocytes exhibit regional heterogeneity across the dorsoventral divide of the spinal cord throughout development and adulthood^21–23^. We expand on these findings by identifying multiple intraspinal regional astrocyte subtypes with distinct molecular profiles that localize to the dorsal, central or ventral grey matter, or the white matter of the healthy adult spinal cord, warranting further research to define their functional contributions. Similarly, astrocyte reactivity is not singular cellular response, but a heterogenous spectrum of context-specific transformations with unique consequences^2,3,5,70,71^. Our results indicate that after SCI, intraspinal regional astrocyte subtypes acquire unique reactivity states, with likely distinct functions and consequences. Reactive LRAs may impact on local circuit function by influencing neurotransmission through region-specific mechanisms. For example, we show that reactive LRAs in the sensory laminae of the dorsal grey matter acquire a late-onset reactivity state (dGM2) with markedly elevated expression of glutamate receptors and transporters. Given the central role of glutamatergic signaling and astrocyte-neuron interactions in neuropathic pain regulation^72^, glutamate sensing and buffering by reactive dGM2 LRAs may play a role in chronic neuropathic pain, a common comorbidity in individuals with SCI. Molecular profiling of the dominant reactive ventral grey matter LRA molecular state in acute iSCI (vGM3) indicates an enhanced capacity for GABA uptake. By limiting extracellular GABA concentrations, vGM3 LRAs may facilitate excitatory neurotransmission that encourages muscle spasticity observed early after injury. In contrast, by sub-acute timepoints, vGM2 astrocytes prevail in the ventral grey matter and exhibit a relatively attenuated capacity for GABA uptake. This may increase extrasynaptic GABA concentrations to promote balanced excitatory-inhibitory neurotransmission and help normalize hindlimb muscle tone to aid recovery of locomotor behavior^73^.

Additionally, reactive LRAs may act through region-specific mechanisms of synaptic remodeling to modulate local circuit function. For example, dGM2 and vGM3 LRAs exhibit elevated expression of distinct synaptogenic proteins, which may facilitate context-specific excitatory synaptogenesis underlying sensory and motor circuit repair after SCI^25,26,74^. vGM2/3 LRAs may also rely on modulation of GABA receptor expression to influence excitatory synaptogenesis via altered production of synaptogenic factors^75^. Moreover, vGM2/3 LRAs express relatively high levels of complement C1q, while dGM2 upregulate Mertk and Megf10 phagocytic receptors, suggesting that reactive LRAs mediate synapse elimination via region-specific mechanisms^76,77^. Interrogation of the astrocyte-intrinsic and extrinsic factors underlying the use of divergent synaptic circuit modulation mechanisms after CNS injury is strongly warranted. Overall, the exploration of how regionally-restricted LRA reactivity states affect local circuit structure, function, and excitability is essential to shaping next-generation treatments that manipulate spared regions of the injured CNS to promote neural repair and recovery.

White matter astrocyte reactivity is remarkably understudied. We identified a type of regionally restricted, white matter degeneration-reactive LRAs that exhibit distinctive, rapid, and persistent expression of *Ccn1*. *Ccn1*^+^ astrocytes intimately associate with phagocytic microglia nodules that clear myelin debris from Wallerian degenerating white matter. White matter astrocytes neighboring degenerating myelinated axons express *Ccn1* to regulate local microglia nodule formation, molecular phenotype, and debris clearance ability (Extended Fig. 10). Astrocyte-secreted CCN1 impacts myelin debris clearance in part via regulation of microglia lipid metabolism, namely limiting cholesterol efflux via regulation of ABCA1 cholesterol exporter. Further investigation is needed to assess the mechanistic and functional relationship between astrocyte CCN1 and microglia sterol metabolism, however, both have been shown to potently regulate macrophage inflammatory profile via a common liver X receptor α-dependent pathway^31,59,78^.

Collectively, our findings across multiple mouse models and human disorders show that *CCN1* expression is an evolutionarily conserved astrocyte-derived que induced by local myelin degeneration. Yet, it is unclear whether *CCN1* expression is stimulated by astrocyte sensing of myelin debris directly^79^, or is rather dependent on cell-extrinsic signals from neighboring injury-reactive cells (e.g. microglia). Moreover, it is attractive to consider that a pathological trigger common to numerous divergent CNS disorders (i.e. myelin damage) may drive an astrocyte response or reactivity state with conserved consequences. Indeed, an important question raised by our findings is whether astrocyte CCN1-mediated signaling can be therapeutically harnessed to enhance white matter debris clearance, restrict inflammation and promote white matter repair across a spectrum of CNS disorders and insults.

Our results indicate that after CNS injury, LRAs acquire heterogenous, evolving, and spatially restricted reactivity states that are mediated by microenvironmental context-specific cues. We show that LRAs retain, but modify their interactions with local cell types, and potently govern multicellular processes underlying degeneration-associated inflammation and tissue repair. This work strongly suggests that the manipulation of LRA reactivity states may be a viable path for limiting chronic neuroinflammation, enhancing functionally meaningful regenerative plasticity, and promoting neurological recovery after CNS injury and in disease.

## Supporting information

SuppInfo1

SuppInfo2

SuppInfo3

## Acknowledgements

This work was supported by: the US National Institutes of Health (NIH) R01NS128094, R00NS105915, K99NS105915 (to J.E.B.), F31NS129372 (to K.S.), R35 NS097303 and R01 NS123532 (RD), R01MH128866, U18TR004146, P30 CA023168 and ASPIRE Challenge and Reduction-to-Practice award (to G.C.); the Paralyzed Veterans Research Foundation of America (to J.E.B.); Wings for Life (to J.E.B.); Cedars Sinai Center for Neuroscience and Medicine Postdoctoral Fellowship (to S.M.); American Academy of Neurology Neuroscience Research Fellowship (to S.M.); California Institute for Regenerative Medicine Postdoctoral Scholarship (to S.M.); The United States Department of Defense USAMRAA award W81XWH2010665 through the Peer Reviewed Alzheimer’s Research Program (to G.C.); The Arnold O. Beckman Postdoctoral Fellowship (to C.E.R.);The Purdue University Center for Cancer Research funded by NIH grant P30 CA023168 is also acknowledged. The authors would like to thank Agilent Technologies Inc. for the donation of the Triple Quadrupole LC/MS instrument to the Chopra Laboratory. The authors would like to thank Dr. Deepti Lall for sharing her expertise in microglia isolation and culture. The authors gratefully acknowledge the International Spinal Cord Injury Biobank (ISCIB) for generously providing the human specimens used in this project, the Cedars-Sinai Applied Genomics, Computation, and Translational (AGCT) Core for RNA-Sequencing and the Cedars-Sinai Biobehavioral Research Core for support in conducting behavioral tests.

## Author contributions

S.M., K.B.S. and J.E.B. designed experiments. S.M., K.B.S., T.I., O.S., C.E.R., G.P.M., R.D., J.P., and J.E.B. conducted experiments. S.M., K.B.S., T.I., A.W.S., O.S., A.P., D.L., B.K., I.Y., R.K., C.H.B., P.M., G.P.M., S.R.V.K., G.C. and J.E.B. analyzed data. S.M., K.B.S. and J.E.B. prepared the manuscript. J.E.B. conceived the study, planned and directed the experiments.

## Methods

### Mice

Young adult male and female mice were used between two and four months of age at the time of the experimental procedure. C57BL/6J mice (JAX: 000664) were used for experiments requiring a wild type (WT) background. For RNA sequencing of astrocyte-specific ribosome-associated mRNA mice expressing RiboTag^19^ (JAX: 029977) were crossed to the well-characterized, astrocyte-specific Cre-driver line, mGfap-cre 73.1237 to generate mGfap-cre-RiboTag mice. mGfap-cre-RiboTag mice were also crossed to Stat3-loxP mice^6^ to generate mGfap-cre-Ribotag-Stat3-loxP mice (Stat3cKO).. Astrocyte-conditional *Ccn1* knockout mice were obtained by crossing the well-characterized, astrocyte-specific Cre-driver line, Aldh1l1-CreERT2^52^ JAX: 031008 to the *Ccn1*-LoxP line^53^ (a gift from Dr. Karen Lyons, UCLA) to generate Aldh1l1-CreERT2-*Ccn1*-LoxP mice (*Ccn1*cKO). CreERT2 expression was activated in young adult mice (6-8 week-old) by administering tamoxifen (Sigma, T5648-1G, 20 mg/ml in corn oil) by subcutaneous injection (100mg/kg, once a day) for five days followed by clearance for three weeks so that no residual tamoxifen remained at the time of experiment initiation. All mice were housed in a facility with a 12 h–12 h light–dark cycle and controlled temperature and humidity, and were allowed free access to food and water. All of the experiments were conducted according to protocols approved by the Institutional Animal Care and Use Committee at Cedars Sinai medical center.

### Surgical procedures

All surgeries were performed on male and female young adult mice (8-12 weeks-old) under general anesthesia with isoflurane in oxygen-enriched air using an operating microscope (Zeiss), and rodent stereotaxic apparatus (David Kopf).

*Spinal cord injury (SCI):* Laminectomy of a single vertebra was performed at spinal cord level T12. Incomplete spinal cord injury (iSCI) by unilateral T12 hemisection was performed on the left side of the spinal cord using a microknife (Fine Science Tools). To be included in the study, mice exhibited complete unilateral hindlimb paralysis for the first 3 days following surgery. A T12 crush SCI was made using no. 5 Dumont forceps (Fine Science Tools) with a 0.4 mm spacer and with a tip width of 0.5 mm. T12 crush animals exhibited paralysis in both hind limbs. In each case, animals received the opiate analgesic buprenorphine subcutaneously before surgery and every 12 h for 48 h after injury. Animals were evaluated thereafter blind to genotype and experimental condition. Daily bladder expression was performed for the duration of the study or until voluntary voiding returned.

*Injections of lysolecithin or myelin into the spinal cord.* 500nL of 1% Lysolecithin or 1 mg/mL CFSE-Myelin in PBS was delivered by intervertebral microinjection to the lateral spinal cord white matter at spinal cord level T12 (coordinates: 200 μm M/L, 300 μm D/V). Injections were carried out at 150 nL/min using finely beveled glass micropipettes connected via high-pressure tubing (Kopf) to 10 μl gastight syringes under the control of microinfusion pumps (Harvard Apparatus). Needles were left in place for 6 minutes prior to being slowly retracted. An equal volume of PBS was injected into the contralateral white matter as vehicle control. Mice were euthanized at 3 days post myelin injection and at 3,10, and 25 days post lysolethicin.

*Sciatic nerve injury:* A small incision was made on the left hindlimb and the two heads of the bicep femoris muscle were gently separated to reveal the sciatic nerve. The sciatic nerve was released from the muscle and elevated using forceps. The isolated nerve was then clamped with hemostats for 10 seconds and then replaced under the muscle. Mice were sacrificed 7 days following sciatic nerve crush.

*Saporin injection:* Conjugated saponins were used to degenerate myelinated and unmyelinated fibers as previously described^66^. Briefly, mice were anesthetized and 8μg (10μl of 0.8μg/μl in PBS) of saporin (non-conjugated control), IB4 conjugated saporin (targets unmyelinated fibers) or CTB-conjugated saporin (targets myelinated fibers) was injected subcutaneously into the plantar surface of the left hind paw foot pad using a 30G insulin syringe. Injections of IB4-saporin and CTB-saporin were considered successful if there was local swelling in the treated hind paw for 24-48 hours following injection. Mice were scarified at 14 and 28 days after injection.

### Experimental Autoimmune Encephalomyelitis (EAE) induction and assessment

Active EAE was induced based on ^80^ with modifications. Nine-week-old C57BL/6 mice were immunized subcutaneously in both hind flanks with 100 μg of myelin oligodendrocyte glycoprotein peptide (MOG_35-55_) emulsified in Complete Freund’s adjuvant containing 200 μg of killed mycobacterium tuberculosis H37Ra (Hooke labs) and injected intraperitoneally (i.p.) on days 0 and 2 with 110 ng pertussis toxin. Assessment of EAE was as follows: 0, no disease; 1, decreased tail tone; 2, hind limb weakness; 2.5, partial hindlimb paralysis; 3, complete hind limb paralysis; 4, front and hind limb paralysis; and 5, moribund state. Animals were collected at different stages of disease based on the following pre-defined criteria: Onset = Partial or completely limp tail (score 0.5 – 1) at day 10 +/- 2 days. Peak = near or complete paralysis of hindlimbs with or without forelimb weakness (score 2.5 – 3.5) at day 14 +/- 2 days. Chronic = mice that reached a score of at least 2.5 (limp tail and incomplete paralysis of hindlimbs) no later than day 16 and collected at day 56.

### Myelin Purification and Conjugation

Myelin was purified from adult C57BL/6 mice brains by sequential ultracentrifugation on discontinuous sucrose gradient and hypo-osmotic shock as previously described^33^. Brains were homogenized with a glass Dounce in 10mM HEPES, 5mM EDTA, and 0.32M sucrose. This was layered on 0.85M sucrose in HEPES/EDTA buffer and centrifuged in a SW41 Ti rotor at 24600 rpm for 30 minutes with acceleration and deceleration = 1. The crude myelin fraction was removed from interface, resuspended in ice-cold distilled water, and centrifuged at 8500 rpm for 15 min. This step was repeated 2 more times. The pellet was then dissolved in 0.3M sucrose in HEPES/EDTA buffer and placed on top of 0.85 M sucrose in HEPES/EDTA. All centrifugation/resuspension steps were then repeated. The final pure myelin pellet was resuspended in PBS, quantified using a BCA assay, and resuspended to 1mg/mL and then conjugated to CFSE as previously described^81^. 1 mg/mL myelin was incubated with 50uM CFSE at 37°C for 15 min. It was then washed with 100 mM glycine in PBS at 14,000 rpm for 15 min. Then washed with PBS twice at 14,000 rpm for 15 min each. Pellets were then resuspended to 1 mg/mL in PBS.

### Hindlimb locomotor evaluation

A modified Basso Mouse Scale (BMS) was developed to evaluate the gradual functional recovery of distinct hind limb muscle groups after iSCI, over time, in freely moving mice. We converted the original BMS protocol^64^ of 5 locomotor categories with a maximal score of 9 into 12 locomotor categories (ankle movement, toe movement, knee movement, weight support, paw placement, dorsal stepping, missing steps, paw position on lift-off, paw position on initial contact, coordination, trunk instability and tail tone) with a maximal score of 37. Analysis was performed at days -5, 0,1,2,3,7,14,28,42, 56,70,84 days post iSCI.

### Cold thermoception behaviral evaluation

Hind paw sensitivity to cold stimuli was evaluated using the acetone test^65^. Spontaneous thermoceptive behaviors were monitored for 1 minute after a drop of acetone (∼25 μl) was applied to the plantar surface of left or right hind paw with the aid of a 22G flexible gavage needle attached to a 1 ml syringe. The total duration of acetone-evoked behaviors (i.e. paw withdrawal, biting, licking, scratching) was measured from videos reviewed in slow motion. Analysis was performed at days -5, 7, 28, and 84 days post iSCI.

### Tissue processing, immunohistochemistry and mRNA *in situ* hybridization

Mice were euthanized by barbiturate overdose followed by cardiac perfusion with 4% paraformaldehyde. Spinal cords were removed, post-fixed for 4-8 hours, and cryoprotected in buffered 30% sucrose. Spinal cords were blocked into 5 mm segments centered around the lesion epicenter, embedded in optimal cutting temperature (OTC) medium and stored at -80°C until sectioning. Serial frozen sections of cervical (C8-T4), thoracic (T3-T12) and lumbar (T9-L3) segments (40 μm, transverse) were prepared using a cryostat microtome (Leica) and stored in antifreeze solution (Glycerol, Sucrose and TBS) at -20°C until processed for evaluation by immunofluorescence and/or mRNA in situ hybridization as described^5^. Primary antibodies include: Rat-CD18 (1:100, Invitrogen), Rat-GFAP (1:1000, Thermofisher), Rabbit-GFAP (1:1000, Dako), Goat-IBA1 (1:1000, Abcam), Rabbit-IBA1 (1:1000, Wako), Rabbit-LPL (1:50, Abcam), Rabbit-PLIN2 (1:500, Progen), Goat-SOX9 (1:200, R&D system), Mouse-SMI32 (1:3000, Biolegend), Sheep-TREM2 (1:250, R&D systems), Rabbit-YAP1 (1:200, Protintech). Mouse primary antibodies were visualized using the M.O.M.® (Mouse on Mouse) Immunodetection Kit (Vector Laboratories). Primary antibodies were selected based on validation for fluorescence immunohistochemistry (IHC) analysis in mouse tissue by the manufacturer, and/or by other investigators based on peer-reviewed publications. Fluorescence secondary antibodies were conjugated to Alexa -488, -Cy3 or -Cy5 all from Jackson Immunoresearch Laboratories. Nuclear staining was performed using 4′,6′-diamidino-2-phenylindole dihydrochloride (DAPI; 2 ng ml−1; Molecular Probes). Sections were cover-slipped using ProLong Glass mounting agent (ThermoFisher). When applicable, tissue sections were incubated in Fluromyelin-Green (1:300) or BODIPY dye (1:1000) (ThermoScientific) prior to DAPI incubation.

Florescent *in-situ* hybridization on fixed-frozen mouse spinal cord sections was performed using RNAscope probes and the Multiplex Fluorescent Detection Kit v2 per manufacturer’s instructions (Advanced Cell Diagnostics). Mouse spinal cord sections were permeabilized with Protease IV. Probes used on mouse spinal cord tissue were as follows: *Abca1* (522251), *AldoC* (429531-c2, 429531-c3), *Arex* (541871), *Ak3* (454791), *Boc* (876211), *Ccn1* (429001), *Gfap* (313211-c2, 313211-c3), *Glipr2* (467171), *Gpnmb* (489511), *Igf1* (443901-c2), *Lair* (509151), *Prdm16* (584281) *Scl1a3* (430781), *Thrsp* (1090411). mRNAs of interest were labeled with the following fluorophores (Akoya): Opal 520 (FP1487001KT), Opal 570 (FP1488001KT), Opal 620 (FP1495001KT), Opal 690 (FP1497001KT). Slides were then processed for immunohistochemistry or stained with DAPI before mounting. Human spinal cord tissue was permeabilized with target retrieval reagent and protease plus. Probes used in human tissue were as follows: *CCN1* (4452081), *GFAP* (311801-C2), *SLC1AA3* (461081-C2). Sections were stained with Dapi and mounted with ProLong Glass or Vectashield mounting medium.

### Imaging

Images of tissue sections used for quantitative analyses were collected using an Apotome epifluorescence microscope with structured illumination hardware and deconvolution software (Zeiss). For whole spinal cord *Ccn1* and microglial analysis, we generated 10x tiles of the entire spinal cord at a single z-plane. Microglial quantification was imaged at 20x (Trem2, LPL) with a z-thickness of 1 μm or 40x (FluroMyelin, SMI32, PLIN2, BODIPY, *Gpnmb* and *Abca1*) with a z-thickness of 0.5 μm. Similarly, images of astrocytes with subtype markers and were imaged at 40x with a 0.5 μm z-stack. Representative images for illustrative purposes were imaged on a Leica SP7 Confocal microscope at 20x or 63x

### Image analysis

Imaris image analysis software (v10) was used to generate 3D volumes of surfaces of IBA1^+^ microglial and a marker of interest (e.g. FluroMyelin, SMI32, PLIN2, BODIPY, LPL, TREM2, LPL). Overlap between IBA1 and marker surfaces (≤ 0.5 μm distance) was used to determine the proportion of microglia that were marker-positive. Similarly, overlap of marker-positive surfaces that were within a IBA1 surface (≤0.5 μm: TREM2, LPL, PLIN2, BODIPY, FluroMyelin, SMI32) determined the volume of marker present within microglia. Measurements were normalized to the total volume to IBA1 microglia and measurements were restricted to the spinal cord dorsal white matter unless stated otherwise. For YAP1 analysis, 3D surfaces were generated for all DAPI^+^ nuclei, YAP1, *Gfap/Slc1a3* mRNA and *Ccn1* mRNA. Astrocyte nucelli were determined by setting the overlap volume of DAPI and *Gfap/Slc1a3* to 15. Astrocytes nuclei expressing *Ccn1* were those containing an overlap volume of *Ccn1* greater than 0.16.

Finally, the YAP1 expression within the *Ccn1^+^* and *Ccn1^-^* astrocytes was the volume of YAP1 (<0um) within these *Ccn1*^+^ *or Ccn1*^-^ astrocyte nuclei. *Spatiotemporal analysis of Ccn1^+^ astrocytes and microglial nodules.* Regional quantification of *Ccn1*+ astrocytes and IBA1+ microglial nodules was performed on 10x image tiles of transverse spinal cord sections using the cell counter plugin (Fiji). Transverse sections were only evaluated if they appeared cytoarchitecturally intact with normal appearing white and grey matter anatomy. Initially, 8 anatomical reference points were used to align transverse spinal cord images: central canal; top of the dorsal white matter; bottom of the dorsal white matter; left and right lateral white matter; top and bottom of the central grey matter; the left and right sides of the central grey matter; the top of the dorsal horn grey matter on left and right sides, and the bottom of the ventral horns on left and right sides. For injured samples, the side containing the majority of *Ccn1*^+^ astrocytes or microglial nodules was labeled as left (ipsilesional). Next, *Ccn1*^+^ astrocytes were quantified as *Gfap*/*Slc1a3* containing nuclei that contained at least 3 *Ccn1* mRNA puncta (RNAscope). Similarly, microglial nodules were quantified as closely associated clusters of microglia containing more than 3 microglial nuclei^33^. At least 2 sections were quantified per animal. *Ccn1*+ astrocytes and microglia nodule counts from different tissue sections were aligned to a common coordinate system using a custom python script. First, all reference and cell coordinates were linearly shifted such that the central canal was set at (0,0). The average of each reference point across all sections per spinal region were used to define a template spinal section which was then used to perform non-rigid transformation (ThinPlateSplineShapeTransformer from the OpenCV2 library) of all cell coordinates. For visualization, *Ccn1* astrocyte /microglia counts were spatially binned per section using a 2D-histogram and counts per bin were averaged per animal and then per condition. The resulting cell density per bin was then plotted.

*Quantitative analysis of in situ mRNA hybridization. Quantitation of RNAscope probe* signal (mRNA) in astrocytes and microglia was carried as described^82^. In brief, thresholding of RNAscope probe signal was first carried out (Otsu method: *Ccn1* and *Gfap/Slc1a3*; Triangle method: *Gpnmb* and *Abca1*) and the area of pixels was then quantified within the soma of *Gfap^+^/Slc1a3*^+^ astrocytes, or IBA1^+^ microglia or microglial nodules, respectively. The area of *Gfap/Slc1a3 and Ccn1* were analyzed from the same astrocyte somas, whereas the area of *Gpnmb and Abca1* mRNA was then normalized to the size of the microglial/nodule.

### Fresh spinal cord tissue collection for astrocyte RiboTag RNA-Seq

Spinal cord tissue was isolated for astrocyte RiboTag RNA-Seq as described^5^. In brief, WT (mGfap-cre-RiboTag) and *Stat3*cKO (mGfap-cre-Ribotag-Stat3-loxP) mice were perfused with ice-cold PBS with heparin and spinal cords were dissected out. 3 mm of spinal cord rostral (T9- T11) and caudal (L1-L3) to the lesion epicenter were then rapidly removed, snap-frozen in dry ice and stored at -80°C until processing for RiboTag RNA-Seq. Spinal cords were collected at 3, 7, 14 and 28 dpi and anatomically equivalent regions of spinal cord were isolated from age- and genotype-matched healthy controls.

### Astrocyte ribosome-associated mRNA isolation, RNA-Seq and analysis

Astrocyte ribosome-associated mRNA was isolated using our previously established methods^5^. In brief, fresh frozen spinal cord tissue was homogenized and haemagglutinin (HA) immunoprecipitation was carried out to purify of astrocyte ribosome-associated mRNA. Astrocyte RNA integrity was analyzed using the 2100 Bioanalyzer (Agilent) with the RNA Pico chip, with RIN ≥ 8 for all samples. RNA concentration was determined using the RiboGreen RNA Assay kit (Life Technologies). cDNA was generated from 10 ng of RNA using the Universal plus mRNA-Seq Kit (Nugen). The workflow consisted of poly(A) RNA selection, RNA fragmentation and double-stranded cDNA synthesis using a mixture of random and oligo(dT) priming, followed by end repair to generate blunt ends, adaptor ligation, strand selection and PCR amplification to produce the final library. Multiplexed sequencing was performed using the NovaSeq 6000 sequencer (Illumina) on a NovaSeq S2 flow cell to produce 50 bp paired-end reads. Data quality was assessed using Illumina SAV and demultiplexing was performed using Illumina Bcl2fastq2 v.2.17. Sequences were aligned to the mouse mm10 genome using STAR aligner (v.2.4.0j). Average percent of uniquely mapped reads was 79 (±8.7)%. Read counts were determined using HT-seq (v.0.6.0). At least 4, and in most cases 6 samples were evaluated per experimental condition. Genes not expressed in minimum of 10 samples (5 counts or more) or average FPKM below 0.75 were filtered out from further analysis. Differential expression analysis (DEA) was conducted using the Bioconductor EdgeR package (v.3.6). DEGs were determined using false-discovery rate (FDR) at 5%. To identify co-regulated astrocyte-enriched genes across time after injury, a gene-gene correlation matrix was constructed using genes that were significantly enriched in astrocytes with a LogFC > 1 and FDR *P*≤0.05 at any timepoint. Astrocyte enriched gene expression was identified by comparing astrocyte HA IP-derived ribosome-associated mRNA to whole tissue mRNA (HA-IP input-derived mRNA). Astrocyte vs whole tissue DEA identified 1249 astrocyte-enriched DEGs, which were used as input for a spearman correlation using log_2_FC changes values from iSCI vs healthy DEA and kmeans clustered into 11 gene modules. Genes in each module were used as input into gene ontology(GO) using Enrichr (“GO_Biological Process_2018” database).

### Nuclei isolation

iSCI mice were perfused with ice-cold PBS with heparin at 3, 7 or 28 dpi, spinal cords dissected out and 3 mm of spinal cord rostral (T9-T11) and caudal (L1-L3) to the lesion epicenter were then rapidly removed, snap-frozen in dry ice and stored at -80°C. An anatomically equivalent region of spinal cord (T11-L1) was isolated from age- and genotype-matched healthy controls. Frozen tissue was homogenized in Homogenization buffer (320 mM Sucrose, 0.1mM EDTA, 0.1% IGAPAL, 5 mM CaCl2, 3 mM Mg(Ac)2, 10 mM Tris, Roche Protector RNAse Inhibitor, Complete Roche Protease Inhibitor Version 12, 0.016mM PMSF, 0.166mM β-Mercaptoethanol; pH=7.8). Nuclei were isolated from the homogenate by iodixanol gradient and resuspended in 1% BSA solution before proceeding immediately to 10x single nuceli RNA-sequencing.

### snRNA-Seq

snRNA-Seq was performed using 10x Chromium Next GEMSingle Cell 3 (v3.1) per manufacturer’s instructions. Samples were loaded to capture 10,000 nuclei per sample. During library preparation, the initial cDNA amplification was run for 13 cycles, which was found to be optimal for 10,000 nuclei. Following library preparation, qPCR was run to quantify library concentration and samples were pooled to equivalent concentrations. Initially a shallow sequencing run of the pooled libraries at ∼20% sequencing saturation, the results of which informed library re-pooling in order to normalize nuclei number within the libraries to obtain ∼40,000 reads/cell. Sequencing was performed by NovaSeq (Illumina) at 2x150 reads base pairs (bps) at 150 pM (average reads/sample: Mean: 2.9E8±1.1E8).

### snRNA-Seq data analysis

Output FASTQ files for each sample were aligned with cellranger-v6.0.2 using the mm10 2020- A reference genome for each sample. Cells matching the following criteria were removed from futher analysis: >5% mitochondrial counts, >25000 counts or <500 counts. Genes expressed in fewer than 50 cells were removed from downstream analysis. Scrublet^83^ was used to remove predicted doublets from each sample. Individual sample data was then concatenated, normalized to 1e4 total counts per cell, log transformed, and batch corrected using Harmony^84^. Quality control thresholding resulted in 230,620 cells from 35 samples for downstream analysis. Cell types were identified based on putative marker genes^38,85–87^. DEG testing utlized sc.tl.rank_genes_groups with method=’wilcoxon’ and corr_method= ‘benjamini-hochberg’ for all comparisons. Nichenet^47^ was performed on astrocytes (‘sender’) and ligands were identified by filtering Nichenet candidates for astrocyte subcluster enrichment relative to all other cell types. The relevant receiver cell type was selected based on NMF cell subtype enrichment. Genes enriched in receiver cell subtype were used as gene set of interest. All expressed genes in the receiver cell subtype were used as the background gene set.

### Spatial transcriptomics

Mouse spinal cord spatial transcriptomics was performed by Visium (10x Genomics). iSCI mice were perfused with ice-cold PBS with heparin at 3, 7, or 28 dpi, spinal cords dissected out and rostral and caudal blocks were rapidly embedded in OTC, snap frozen on dry ice and stored at - 80°C until sectioning. Visium slides were pre-chilled in a cryostat (Leica) for 30 minutes at the time of sectioning. Two 10um sections were taken from lesion-remote rostral (T9-T11) and caudal (L1-L3) blocks, equivalent to samples analyzed by snRNA-Seq. Samples were processed using the Visium Spatial Gene Expression Reagent Kit (10X Genomics) as per manufacturer’s established protocol. cDNA libraries were pooled in a NovaSeq6000 SP v1.0 flowcell and paired-end sequencing was performed on an Illumina NovaSeq6000 sequencer.

### Spatial transcriptomics analysis

Spots overlaying tissue sections were manually annotated in the Loupe (10X Genomics) and processed by spaceranger-v1.3.0 and aligned against the mm10 reference genome mm10- 2020-A. Hematoxylin and eosin staining of transverse spinal cord sections was used to manually annotate lateral white, ventral white, dorsal white, central grey, dorsal horn, and ventral horn. Additionally, gene expression of inflammation and gliosis-associated genes was used to distinguish lesion ipsilateral and contralateral sides. Quality control thresholding resulted in 14,566 spots across 16 biological replicates (n=4 mice/group; 2 sections per rostral and caudal block). Data was normalized to 1e4 counts and log transformed before running PCA and UMAP projection. To accommodate for morphological variation, a non-rigid transformation was applied (ThinPlateSplineShapeTransformer from the OpenCV2 library) using manually placed neuroanatomical reference points, in a manner equivalent to aligned average density plot construction for *Ccn1*^+^ astrocyte and WDM nodules counts. Tissue alignment was validated by examination of known spatially restricted gene expression (e.g. Mbp,Syp). Cell2location was used to spatially integrate snRNAseq subclusters and spatial transcriptomics data. The top 30 highest expressed genes from each dataset, mitochondrial genes, and genes expressed in <5% of cells/spots and with mean<1.12 were filtered out to generate snRNA-seq input. Cell2location was run with the following parameters: batch_key= “Date library prep”, continuous_covariates=”total_counts”, categorical_covariates=”User”, N_cells_per_location=12, detection_alpha=200) and trained for 40,000 epochs. The cell2location matrix was used as input for NMF to identify spatially co-occurring cell types. NMF from sklearn was run with the following parameters: (n_components=8, alpha=0.9,max_iter=1000, shuffle=True, init=”nndsvda”,l1_ratio=0.9).

### Microglial isolation (lipidomic and culture)

Mice were perfused with ice-cold PBS with heparin and the brain or spinal cord was freshly dissected. For iSCI mice, 1mm rostral and caudal to the lesion epicenter was removed and discarded, and the entire injured lateral side of the spinal cord (rostral and caudal to the lesion) was collected for microglial isolation. Dissected tissue was chopped with a sterile razor and then dissociated using the Neural Tissue Dissociation Kit (P) (Miltenyi Biotech) and the GentleMACS dissociator with heaters per the manufacturer’s instructions. Following dissociation, samples were filtered through a 70μm strainer, and myelin was depleted using Myelin removal beads II (Miltenyi Biotech) using the AutoMACS separator. Finally, microglia were isolated by incubating samples with CD11b microbeads (Milteyni) and isolated with the AutoMACS as per the manufacturer’s instructions. Cell number was then determined before proceeding to downstream applications (i.e. lipidomics, culture).

### Microglial isolation (lipidomic and culture)

Mice were perfused with ice-cold PBS with heparin and the brain and spinal cord were freshly dissected following. For iSCI mice, 1mm rostral and caudal to the lesion epicenter was removed and discarded, and the injured lateral side of the spinal cord (rostral and caudal to the lesion) was collected for microglial isolation. Dissected tissue was minced with a sterile razor blade and then dissociated using the Neural Tissue Dissociation Kit (P) (Miltenyi Biotech) and the GentelMACS dissociator with heaters per the manufacturer’s instructions. Following dissociation, samples were filtered through a 70μm strainer, and myelin depleted using Myelin removal beads II (Miltenyi Biotech) and the AutoMACS separator. Finally, microglia were isolated by incubating samples with CD11b microbeads (Milteyni) and isolating with the AutoMACS as per the manufacturer’s instructions. Cell number was then determined before proceeding to downstream applications (i.e. lipidomics, culture).

### Sample Preparation for Lipid Extraction

Lipid extraction from the frozen microglial cell pellets was done following Bligh and Dyer protocol with slight modifications^88,89^. Briefly, the pellets were thawed at 4 °C for 10 mins, after which, cold methanol, and HPLC-grade chloroform were added in a ratio of 2:1. The samples were vortexed for 10 seconds, resulting in a one-phase solution which was then incubated for 15 mins at 4°C. A biphasic solution was then obtained by adding ultrapure water and chloroform in a 1:1 ratio. Next, the samples were centrifuged at 16,000 x g for 10 mins, giving rise to 3 phases in each tube. The bottom phase in the tube is the organic phase that contains lipids. Next, the solvents from the organic phase were evaporated using SpeedVac vacuum concentrator for 1 hour resulting in dried lipid extracts.

### Unbiased Lipidomics using Multiple Reaction Monitoring (MRM)-Profiling

Dried lipid extracts from microglial cells were reconstituted in 200 µL of a 50:50 methanol:chloroform solution containing 10 mM ammonium formate. Prior to analysis, lipid extracts were further diluted in acetonitrile: methanol:70:30, with 10 mM ammonium formate. The “quality control” sample was the injection solvent containing 0.02 µg/mL of the quantitative mass spectrometry internal standard EquiSPLASH® (Avanti Polar Lipids, #330731), which was monitored over time to ensure the instrument’s appropriate operation. All MRM profiling experiments were conducted on an Agilent 6495C triple quadrupole mass spectrometer (Santa Clara, CA) outfitted with an Agilent 1290 Infinity II LC system and G7167B autosampler (Santa Clara, CA). A volume of 8μL of diluted lipid extract was introduced into the Agilent Jet Stream (AJS) ion source of the mass spectrometer by flow injection for each MRM method. Briefly, MRM methods were established for 10 lipid classes and spanned 1497 individual lipid species^89,90^. Lipid classes of interest were phosphatidylcholine (PC), phosphatidylethanolamine (PE), phosphatidylglycerol (PG), phosphatidylinositol (PI), phosphatidylserine (PS), acyl carnitine (CAR), cholesterol ester (CE), diacylglycerol (DG), triacylglycerol (TG), and sphingomyelin (SM).

### Lipidomics Data Analysis

Statistical analysis of MRM transitions in lipids for all sample comparisons were conducted using the edgeR software package^91^ based on our previous study^89,90^. The ion count of each lipid was denoted by ‘l’ for a given sample ‘s’. An ‘intercept’ sample, representing the experimental blank (injection medium), was included to ensure that all comparisons are meaningful relative to the blank. The edgeR package^3^ fits a generalized linear model to a log-linear formula for mean variance relationship as follows:

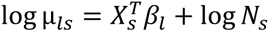

This formula calculates the total ion intensity for each sample ‘s’, summing to ‘N_s_’. This approach enables the determination of the coefficient of variation (CV) for the ion count of each lipid in a sample (‘*y_ls_*’). The dispersion (Φ_l_) of each lipid and is calculated using the common dispersion method.^5^ Based on these values, the log_2_ fold change (log_2_FC) between samples is calculated, and the corresponding P values are derived using the likelihood ratio test. P values less than 0.05 were considered significant.

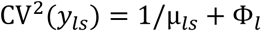

### Microglial culture and cholesterol efflux assay

Primary microglia isolated from male and female mice aged (8-12 weeks) were seeded at 1x10^5^ cells/well in a flat-bottom 96-well plate coated with poly-L-lysine (0.01%, Sigma-Aldrich). Cells were grown in microglia media (10% fetal bovine serum, 1x penicillin-streptomycin, 10ng/mL carrier-free (CF) recombinant mouse GM-CSF, 10ng/mL CF recombinant mouse M-CSF, 10ng/mL CF recombinant human TGFβ1, in DMEM/F-12 Ham) for 4 days at 37°C, 5% CO_2_ with full media change after 24 hours. Microglia cholesterol efflux was assessed using a Fluorometric Cholesterol Efflux Assay Kit (Abcam; ab196985) following the manufacturer’s instructions. Briefly, on day 4 microglia were loaded with fluorescent cholesterol for one hour, then placed into equilibration media containing CCN1 (50ng/ml in 0.1%BSA) or Vehicle (0.1% BSA) for 16 hours. Following incubation, cells were washed with phenol red-free DMEM/F-12 Ham and incubated with cholesterol acceptor solution (2% (2-Hydroxypropyl)-β-cyclodextrin) for 6 hours. Plates were then centrifuged for 2 minutes at 1000 x g and the cell supernatant was collected for fluorometric analysis of cholesterol content. Meanwhile, adherent cells were lysed and processed for fluorometric cholesterol content within the cell. Fluorometric measurements were performed using Varioskan LUX (ThermoFisher) at (Ex/Em = 485/523 nm). Percent cholesterol efflux was then calculated for each sample by dividing relative fluorescence unit (RFU) of the supernatant by the total cholesterol content (RFU of supernatant + cell lysate). The effect of treatment was then calculated by subtracting the percent cholesterol efflux of the negative control (no cholesterol acceptor) from the percent cholesterol efflux of treatment (CCN1 or vehicle), followed by normalization with the percent cholesterol efflux of vehicle.

### Human MS and SCI spinal cord tissue

Human formalin-fixed paraffin-embedded (FFPE) spinal cord tissues from individuals with MS and neurologically healthy controls were prepared from autopsy-derived tissues collected by the rapid autopsy protocol approved by the Cleveland Clinic Institutional Review Board. Transverse spinal cord sections (7 μm) were prepared and the demyelinated lesions were identified by loss of proteolipid protein (PLP) immunoreactivity. FFPE spinal cord tissues from individuals with SCI and associated clinical and neuropathological information were obtained from the International Spinal Cord Injury Biobank (ISCIB; Vancouver, Canada). The Clinical Research Ethics Board of the University of Columbia (Vancouver, Canada) granted permission for post-mortem spinal cord acquisition and for sharing biospecimens. Spinal cord biospecimens were collected from consented participants or their next-of-kin and provided as FFPE tissue sections at a thickness of 5 μm. SCI tissue sections evaluated herein derive from lesion-remote regions of the injured cord that exhibit white matter damage and/or Wallerian degeneration as determined by an experienced ISCIB neuropathologist based on combined LFB with H&E staining, and results from 7-Tesla magnetic resonance imaging of these spinal cord tissue blocks prior to sectioning. Deidentified information for healthy, MS and SCI patients is provided in Extended data figure 9m.

### Statistics and reproducibility

Statistical evaluations of repeated measures were performed using one-way or two-way ANOVA with post hoc independent pairwise analysis using Tukey tests, Wilcoxon rank sum test, or t-tests using Prism 8 (GraphPad). *P* values are reported in the figures or figure legends. Differences with P < 0.05 were considered to be statistically significant. Power calculations were performed using G*Power Software v.3.1.9.2. All immunohistochemistry, *in situ* hybridization, and *in-vitro* culture experiments analyses shown were repeated at least three times with similar results. Experimental procedures and quantitative analyses were conducted by individuals blinded to experimental group assignments.

### Data availability

Raw and normalized genomic data have been deposited at the NCBI Gene Expression Omnibus under the SuperSeries accession number GSEXXXXXX.

---

**Extended Data Fig. 1.**
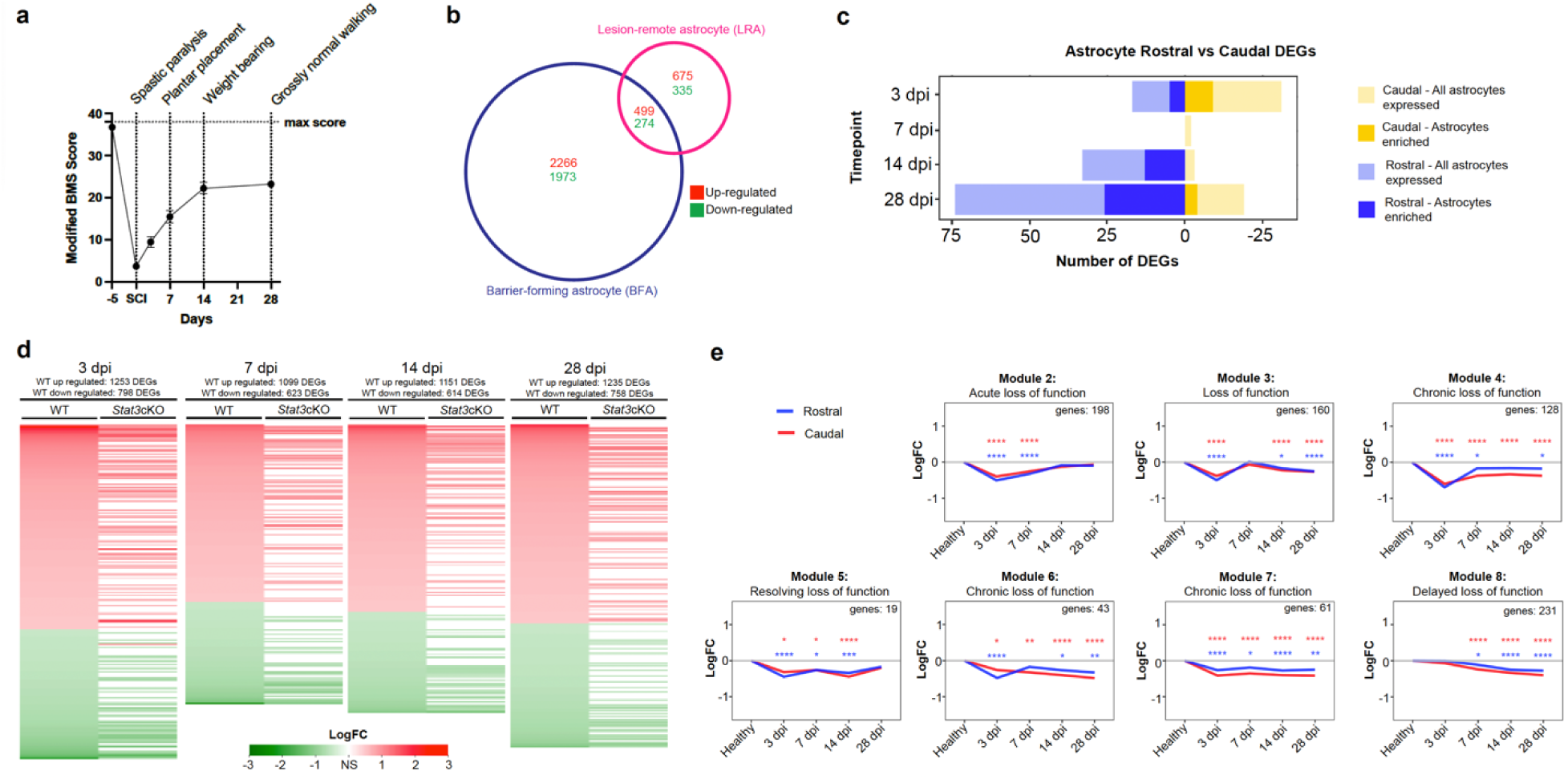
**a,** Spontaneous recovery of locomotor behavior in the left hindlimb after iSCI as scored by a modified BMS protocol. **b**, Venn diagram showing DEGs in 14 dpi BFAs vs LRA, relative to healthy spinal cord astrocytes (FDR *P* ≤ 0.01)**. c**, Bar graph illustrating rostral and caudal LRA RiboTag RNA-Seq DEGs over time after iSCI, vs Healthy (FDR *P* ≤ 0.01). **d**, Heatmaps illustrating *Stat3*cKO effects on WT LRA RiboTag RNA-Seq DEGs over time after iSCI, (vs Healthy; rostral and caudal combined; FDR *P* ≤ 0.05, Log2FC ≥ 0.5). **e**, Line graphs illustrating temporally regulated gene expression of LRA co-regulated gene modules rostral and caudal to the injury (\**P* ≤ 0.05, \*\**P* ≤ 0.002, \*\*\**P* ≤ 0.0002, \*\*\*\**P* ≤ 0.0001, Two-way ANOVA with Tukeys). Lines represent average expression across all co-regulated genes.

**Extended Data Fig. 2.**
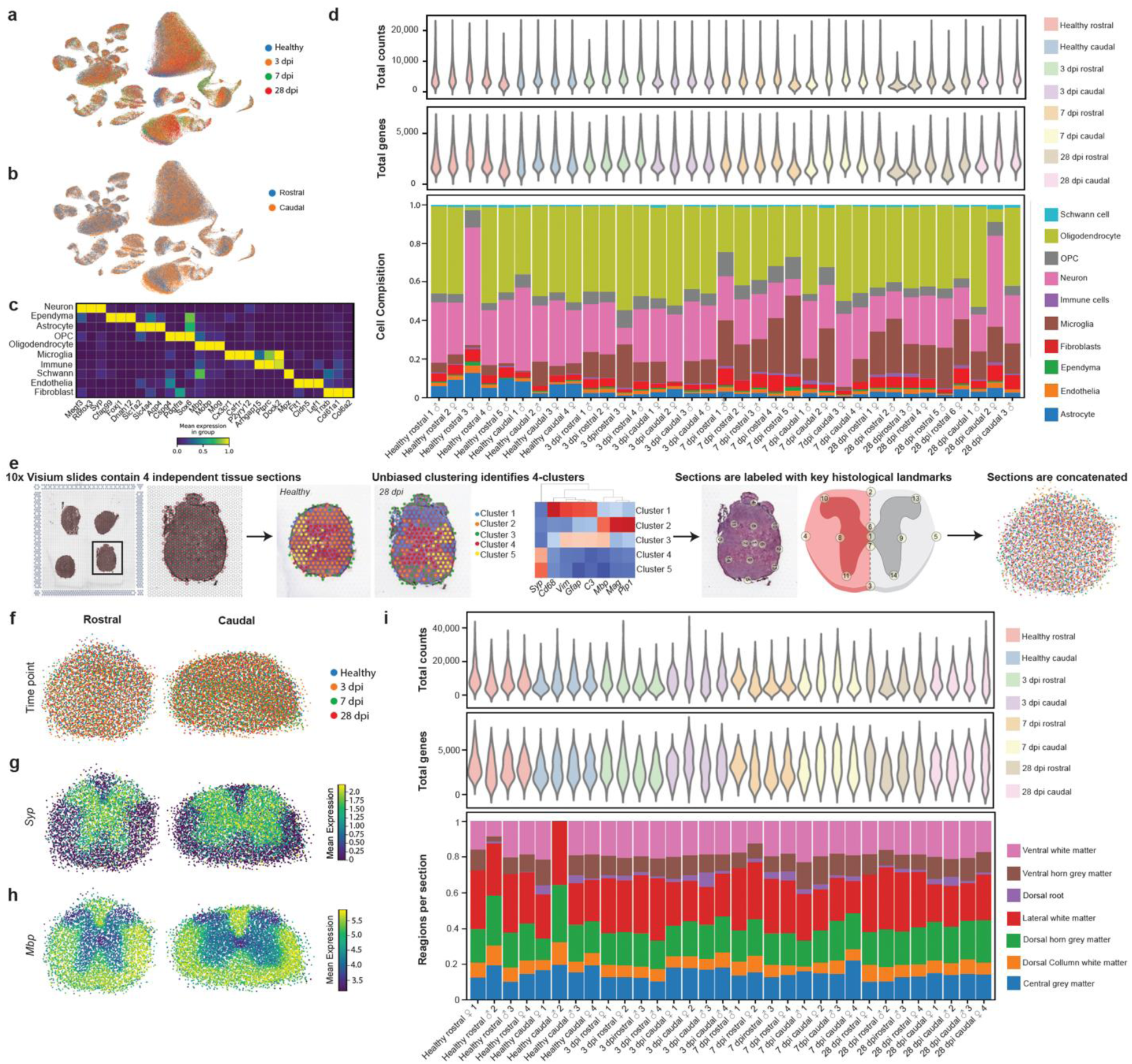
**a**, UMAP of the final snRNAseq dataset after quality control (230,570 nuclei) colored by timepoint (healthy, 3dpi, 7dpi, 28dpi) and **b**, spinal region (Rostral vs Caudal). **c**, Heatmap of putative marker genes used to identify different cell types. **d**, Violin plots showing the distribution of total counts and genes detected, and a stacked bar plot showing the proportions of broad cell types for each individual snRNAseq sample. All samples showed similar quality control statistics and cell type contribution. **e**, Schematic showing pipeline for intraspinal regional spatial transcriptomics. For iSCI tissue sections, unbiased clustering identifies the lesion ipsilateral spinal cord white matter, which exhibits elevated expression of inflammation and gliosis genes relative to contra-lesional spinal cord regions. **f**, Aligned Visium data shown for the rostral and caudal regions in space and colored by timepoint (healthy, 3dpi, 7dpi, 28dpi). **g**, **h**, Mean expression for Synaptophysin *(Syp)* and Myelin Basic Protein (*Mbp)* confirming localization to the grey and white matter respectively. **i**, Violin plots showing the distribution of total counts and genes detected, and a stacked barplot showing the proportions of spinal region for each individual Visium experimental replicate. All replicates showed similar quality control metrics and tissue region contribution.

**Extended Data Fig. 3.**
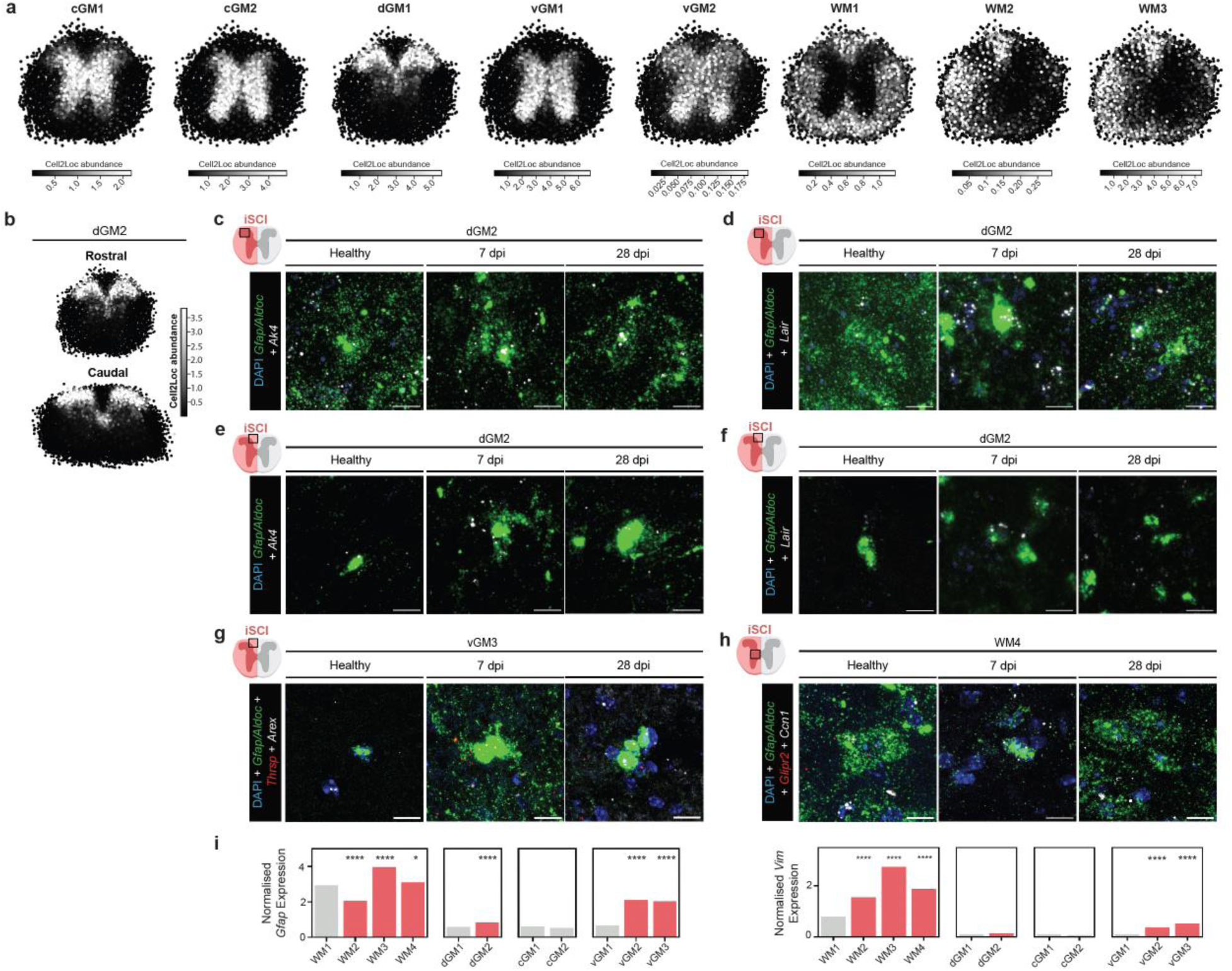
**a**, Spatial characterization (Cell2Location abundance plots) of regionally-restricted astrocyte molecular states across lesion-remote rostral spinal cord. Astrocyte cluster 11 (not shown) did not exhibit a clear anatomical spatial patter, attributable to the small number of cells in this cluster. **b**, Spatial characterization (Cell2Location abundance plots) of dGM2 astrocytes in lesion-remote rostral and caudal spinal cord. **c-f**, High magnification of dGM2 marker *Ak4 and Lair* shows elevated expression in astrocytes *(Gfap^+^*/*Aldoc^+^*) in dorsal horn grey matter, but not white matter. **g**, High magnification of vGM2 markers *Thrsp* and *Arex* shows lack of injury-reactive astrocyte expression in white matter. **h**, High magnification of WM4 markers *Glipr2* and *Ccn1* show lack of injury-reactive astrocyte expression in grey matter. **i**, Mean expression of *Gfap* and *Vim* is used to identify hypertrophy-associated astrocyte molecular states (red bars) detected by snRNA-Seq expression. Differential expression testing for *Gfap* and *Vim* determined relative to regional healthy cluster by Wilcoxon rank sum test. \**P* ≤ 0.05, \*\*\*\**P* ≤ 0.0001. Scale bars: 10 µm.

**Extended Data Fig. 4.**
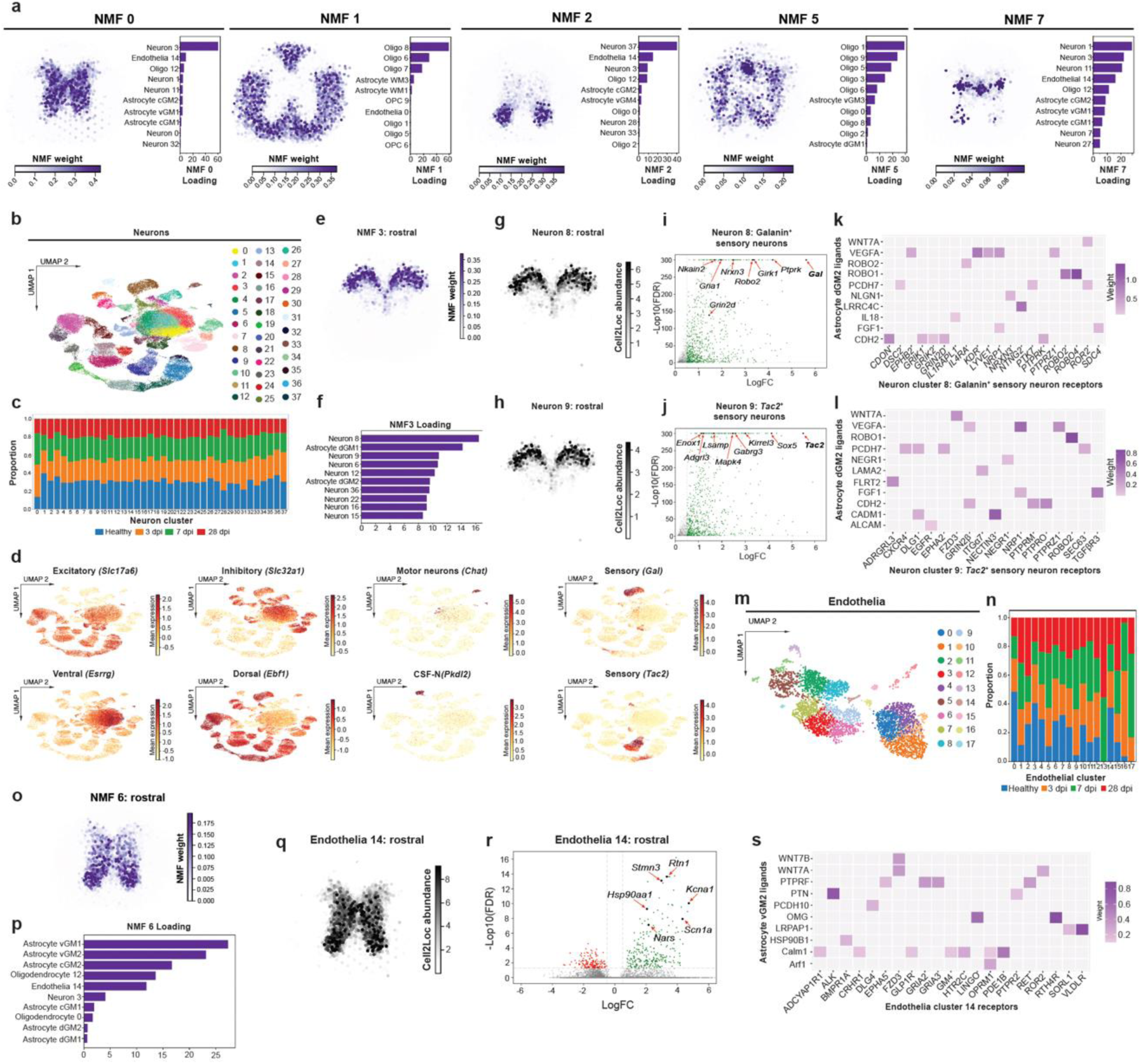
**a**, Spatial profiles of additional NMF factors. **b**, UMAP of spinal cord neuron subtypes identified by snRNA-Seq for healthy and all post-injury time points, rostral and caudal. **c**, Neuron subtype proportions across healthy and iSCI reveals little injury-reactive alterations in neuron subtype representation after iSCI. **d**, UMAP of spinal cord neurons reflecting expression of established subtype markers. **e-h**, Spatial and cell identity loading profiles of NMF3 revealed that dorsal horn astrocytes (dGM1,dGM2) intermingle with multiple subtypes of sensory neurons of the superficial laminae (Neuron 8, 9). **i**, **j**, Volcano plots of DEGs in Neuron 8 *Gal*^+^-expressing inhibitory interneurons or Neuron 9 *Tac2^+^* excitatory interneurons, relative to other neuron subtypes (FDR *P* ≤ 0.01). **k**, **l**, NichNet analysis identified putative ligands from reactive dGM2 astrocytes (senders) and predicted receptors enriched in Neuron 8 or Neuron 9 (receivers) (NicheNet). **m**, UMAP of spinal cord endothelial cell subtypes identified by snRNA-Seq for all time points, rostral and caudal. **n**, Endothelia subtype proportions across healthy and iSCI groups reveals multiple injury-reactive alterations in endothelia subtype representation after iSCI. **o-q** Spatial and cell identity loading profiles of NMF6 revealed that ventral grey matter reactive LRAs (vGM2) intermingle with local endothelia 14. **r**, Volcano plot of Endothelia 14 DEGs, relative to other endothelia subtypes (FDR *P* ≤ 0.01). **s**, NichNet analysis identified putative ligands from reactive vGM2 astrocytes (senders) and predicted receptors enriched in Endothelia 14 (receivers).

**Extended Data Fig. 5.**
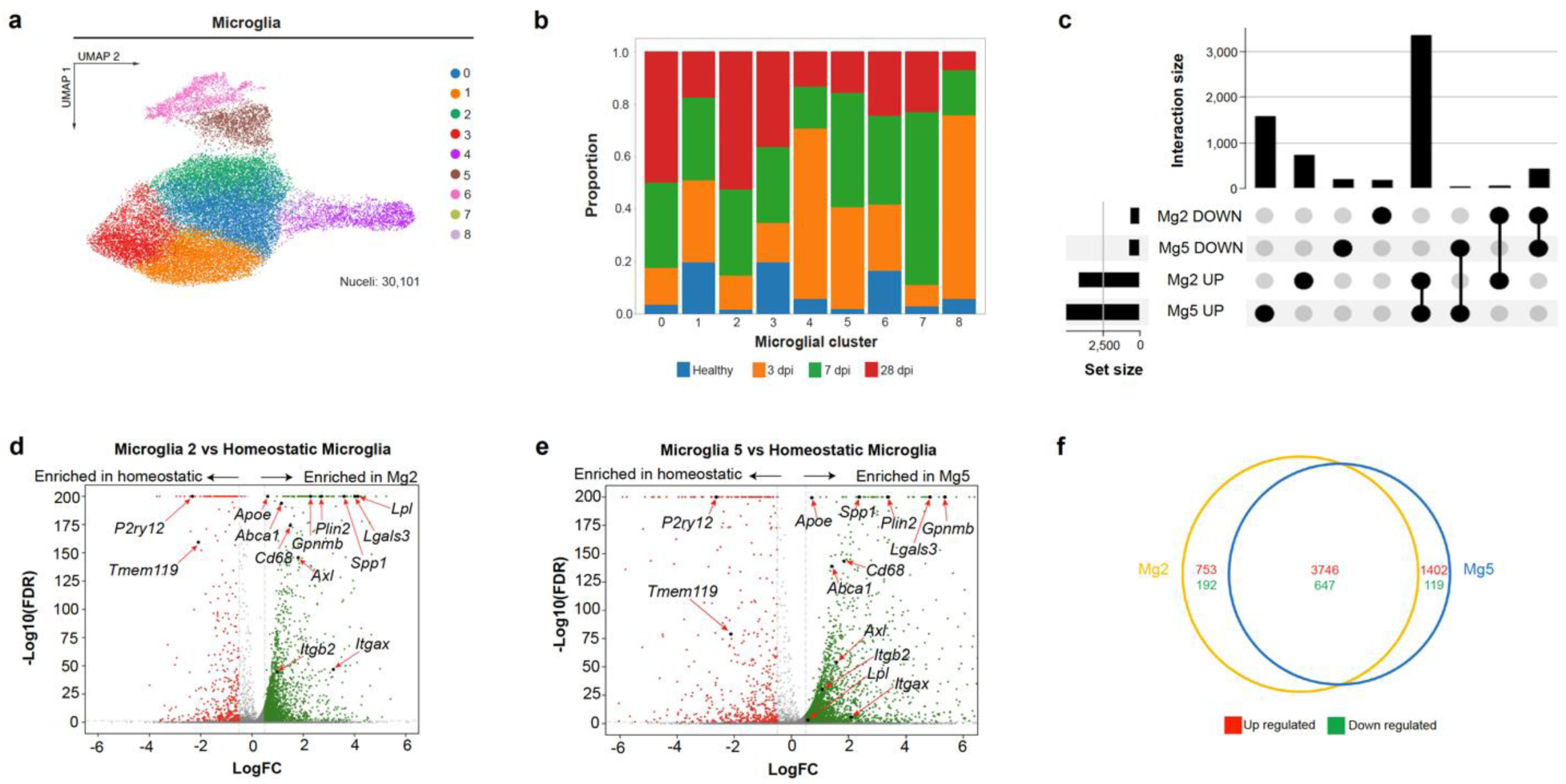
**a**, UMAP of spinal cord microglia subtypes identified by snRNA-Seq for healthy and all post-injury time points, rostral and caudal. **b**, Microglia subtype proportions across healthy and iSCI groups reveals multiple injury-reactive alterations in subtype representation after iSCI. **c**, Upset plot of Mg2 and Mg5 microglia DEGs relative to all microglia subtypes. **d**, **e**, Volcano plots of Mg2 and Mg5 microglia DEGs relative to homeostatic microglia subtype. **f**, Venn diagram comparing Mg2 and Mg5 microglia DEGs relative to homeostatic microglia subtype. DEG FDR *P* ≤ 0.01.

**Extended Data Fig. 6.**
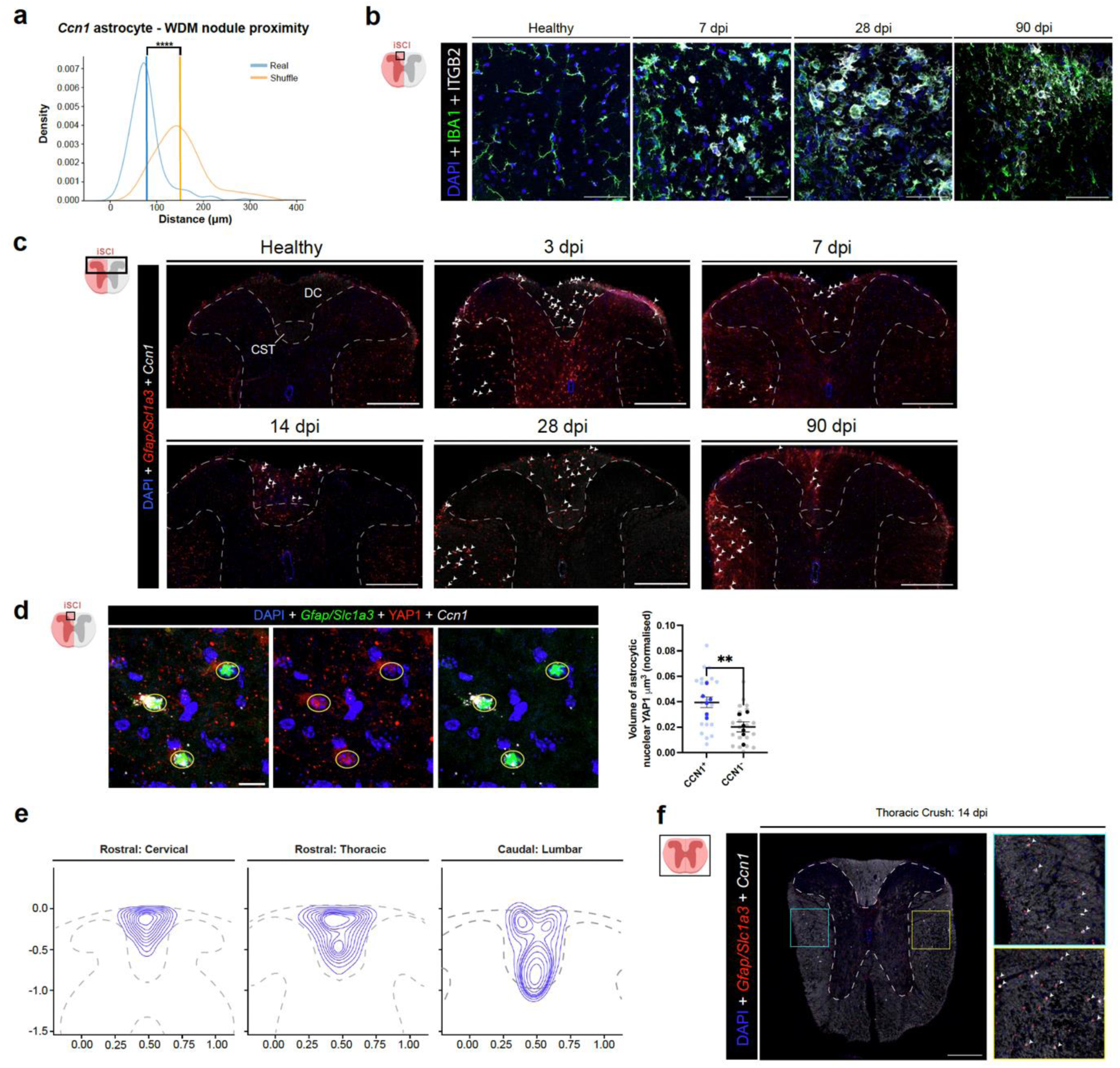
**a**, Plotting the distribution of distances of *Ccn1*^+^ astrocytes to WDM nodules within Wallerian degenerating dorsal column white matter relative to a randomly shuffled distribution illustrates that WDM nodules are more likely to be proximate to *Ccn1*^+^ astrocytes (mean=78μm) than would be expected by random chance (mean = 150µm). ****=*P*<0.001, Wilcoxon test. **b**, ITGB2 levels in IBA1^+^ microglia healthy and over time after iSCI. **c**, Low magnification of *Ccn1*^+^ astrocytes (white arrowheads) in healthy vs lesion-remote spinal cord (rostral). **d**, Comparison of nuclear YAP levels in *Ccn1*^+^ and *Ccn1*^-^ astrocytes (*Gfap*^+^/*Slc1a3*^+^) (n=6, \*\**P* ≤ 0.005, Nested t-test). Graph shows mean ± SEM. Opaque and transparent data points illustrate experimental replicate mean and counts from individual cells, respectively. **e**, Drawing the 2-D kernel density estimate of *Ccn1*^+^ astrocytes within the dorsal white matter at different spinal cord levels illustrates that their spatial distribution closely follows that of Wallerian Degeneration after iSCI. Rostral to the lesion (cervical, thoracic), *Ccn1*-expressing astrocytes localize to the Wallerian degenerating DC. Caudal to the lesion, *Ccn1*-expressing astrocytes localize mainly to the Wallerian degenerating corticospinal tract. **f**, *Ccn1*^+^ astrocytes (white arrowheads) are observed bilaterally after crush SCI that damages both sides of the spinal cord. Scale bar: b, d=50 µm c,f: 250 µm

**Extended Data Fig. 7.**
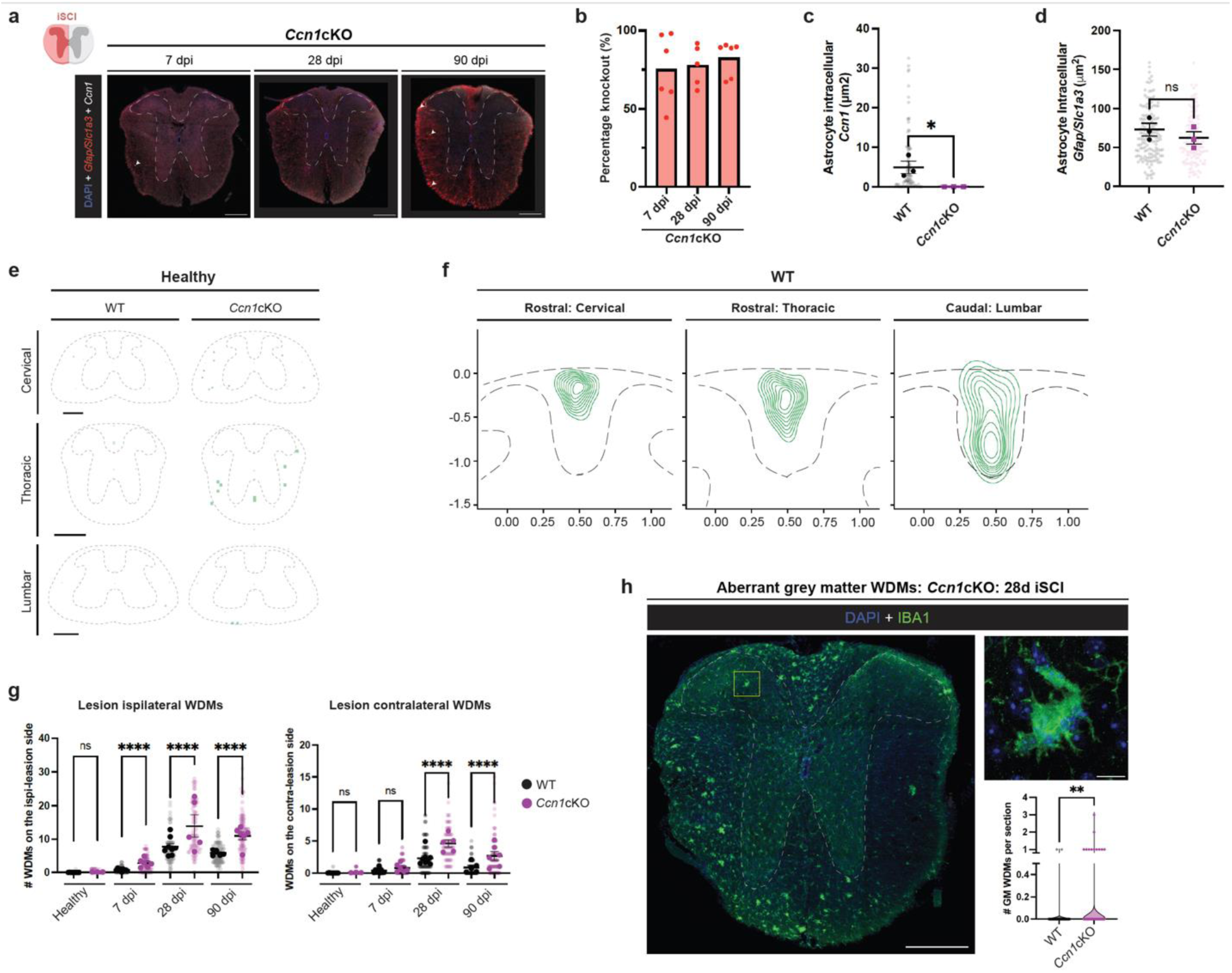
**a**, Low magnification of *Ccn1* and *Gfap/Slc1a3 (arrowheads)* in iSCI *Ccn1*cKO spinal cord. **b**, Percentage *Ccn1* knockout as proportion of WT *Ccn1*^+^ astrocytes. **c**, **d**, Quantification of *Ccn1* and *Gfap/Slc1a3* expression in individual astrocytes shows reduction in *Ccn1* expression per astrocyte, while *Gfap/Slc1a3 r*emain equivalent to WT astrocytes (n=3 mice/genotype, 23-53 astrocytes analyzed/mouse; *P ≤ 0.05, Student’s t-test). **e**, Aligned average density plots of WDM nodules in healthy WT and *Ccn1*cKO spinal cord. **f**, Spatial characterization of *WDM nodules* relative to Wallerian degenerating DC (cervical, thoracic) and CST (lumbar). **g**, Quantification of WDM nodules in the ipsi-lesion and contralateral white matter after iSCI in WT vs *Ccn1*cKO (\*\*\**P* ≤ 0.0002, \*\*\*\**P* ≤ 1x10^-4^, two-way ANOVA with Tukeys). **h**, Low and high magnification of IBA1^+^ WDM nodules in *Ccn1*cKO white matter at 28 dpi. **i**, Comparison of grey matter WDM counts per spinal cord section in iSCI WT vs *Ccn1*cKO (346-411 sections analyzed/genotype; \*\**P* ≤ 2x10^-3^, Student’s t-test). Graphs show mean ± SEM. Opaque and transparent data points illustrate experimental replicate mean and counts from individual tissue sections, respectively. Scale bar a,e,h: 250 µm, h-inset 10 µm

**Extended Data Fig. 8.**
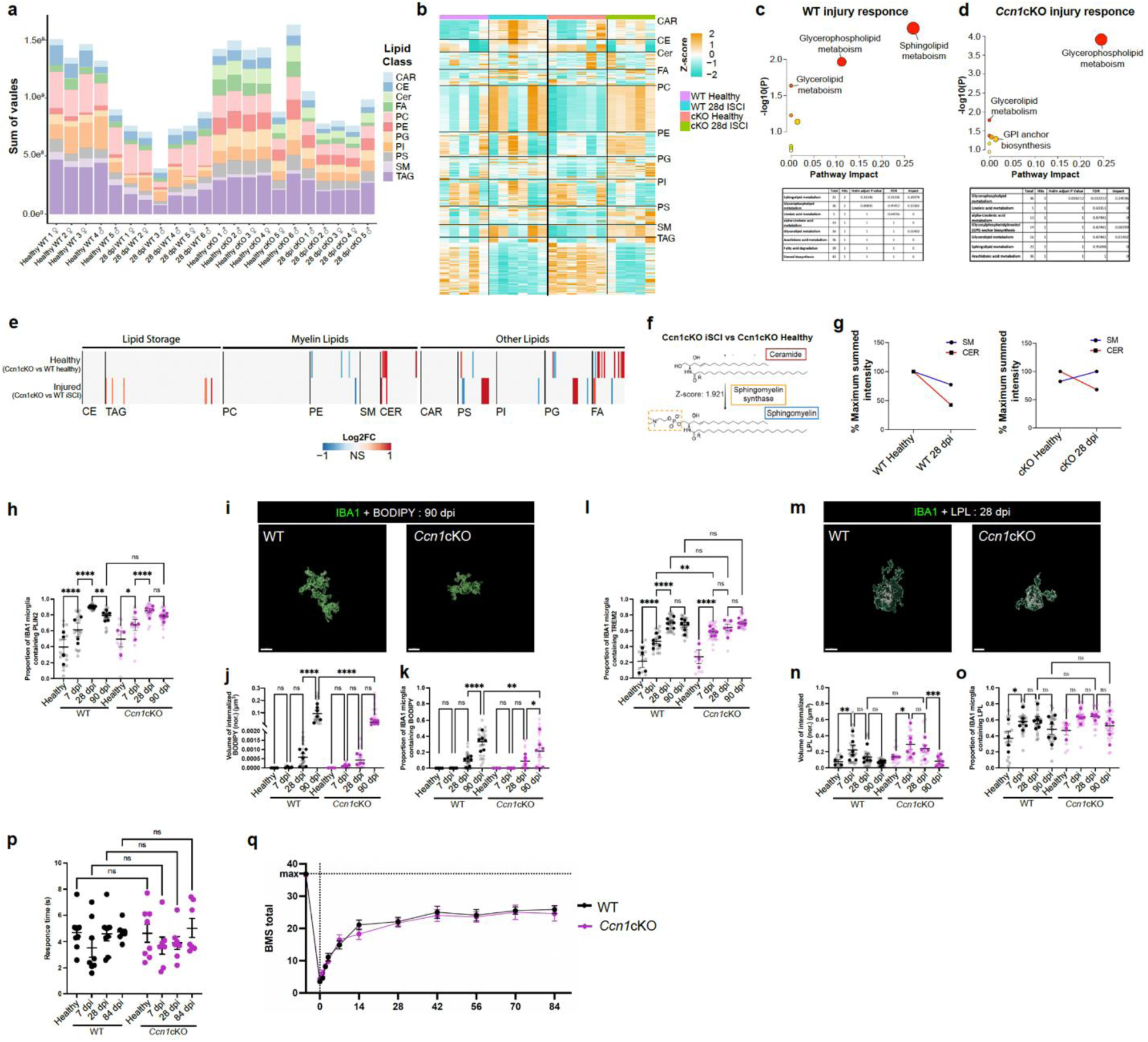
**a**, Microglia lipid species distribution across treatment groups. Note that in both WT and *Ccn1*cKO animals, the total microglia lipidome is reduced at 28 dpi. **b**, Heat map of Z-scores showing relative levels of all significant lipid subtypes detected by unbiased microglia lipidomic analysis**. c**, **d**, Lipid pathway enrichment of the WT and *Ccn1*cKO injury response highlighting a shift in the predominant lipid pathways employed after iSCI. **e**, Direct pairwise comparison of WT and *Ccn1*cKO microglia lipid profile for healthy and iSCI, including lipid droplet- and myelin-associated lipid subtypes (Log_2_ fold-change FDP *P* ≤ 0.01). **f**, Schematic of ceramide to sphingomyelin conversion mediated by sphingomyelin synthase predicted by Biopan lipid pathway analysis comparing the iSCI and healthy *Ccn1*cKO animals (z-score 1.921). **g**, Plots show percentage maximum summed intensity for sphingomyelin (SM) and ceramide (CER) in both WT and *Ccn1*cKO mice. **h**, Quantification of the proportion of PLIN2 containing microglia from WT or *Ccn1*cKO *Ccn1*cKO Wallerian degenerating dorsal column white matter. **i-k**, High magnification 3D image and quantitative comparison of BODIPY^+^ lipid droplets within IBA1^+^ WDM nodules from WT and *Ccn1*cKO Wallerian degenerating dorsal column white matter (n=3-6 mice/group). **l**, Relative proportions of IBA1^+^ microglia that are TREM2^+^ from WT and *Ccn1*cKO Wallerian degenerating dorsal column white matter. **m-o**, High magnification and 3D image and quantitative comparisons of IBA1^+^ microglia that contain LPL (n=3-6 mice/group). **p**, Line graph illustrating spontaneous recovery of locomotor behavior in the left hindlimb after iSCI in WT vs *Ccn1*cKO mice by modified Basso mouse scale scoring. Graphs show mean ± SEM. Opaque and transparent data points illustrate experimental replicate mean and counts from individual tissue sections/cells, respectively. Unless stated otherwise, \**P* ≤ 0.05, \*\**P* ≤ 0.002, \*\*\**P* ≤ 0.0002, \*\*\*\**P* ≤ 0.0001, Two-way ANOVA with Tukeys. Scale bars, 10 µm

**Extended Data Fig. 9.**
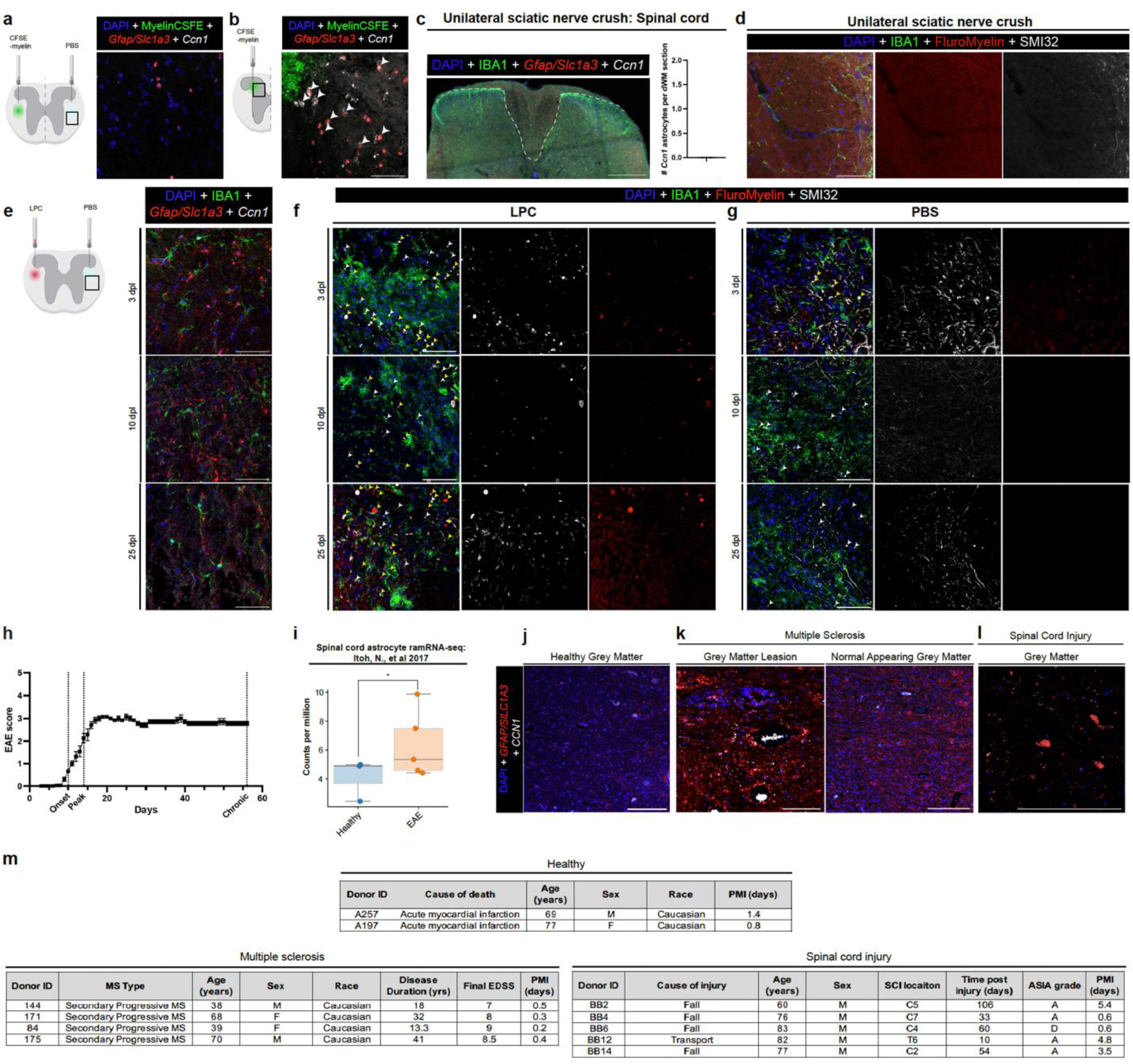
**a**, High magnification of *Ccn1* expression labeling and astrocytes (*Gfap*^+^/*Slac1a3*^+^ cells) in spinal cord lateral white matter 72 hrs following injection of PBS (CFSE-myelin vehicle control). **b,** High magnification of *Ccn1^+^* astrocytes (*Gfap*^+^/*Slac1a3*^+^ cells, white arrowheads) in spinal cord grey matter following injection of CFSE-conjugated myelin into dorsal horn grey matter. **c**, Low magnification and quantification of *Ccn1^+^* astrocytes (*Gfap*^+^/*Slac1a3*^+^ cells) in dorsal half of lumbar spinal cord following unilateral (left) sciatic nerve crush. **d**, Evaluation of myelin and axon degeneration in spinal cord dorsal white matter following sciatic nerve crush. **e**, *Ccn1* expression and astrocytes (*Gfap*^+^/*Slac1a3*^+^ cells) in spinal cord lateral white matter following injection of PBS (LPC vehicle control). **f-g**, Assessment of myelin degeneration (Fluoromyelin, yellow arrowheads) and damaged axons (SMI32, white arrowheads) in spinal cord lateral white matter following demyelination by LPC or injection of PBS. **h**, EAE disability severity scores assessing locomotor disability (n=4-7 mice/group). Graph shows mean ± SEM across experimental replicates. **i**, Astrocyte RiboTag RNA-Seq from mouse chronic EAE spinal cord shows significantly elevated *Ccn1* expression^68^ (FDR P ≤ 0.01). Graph show mean ± SEM. Data points illustrate experimental replicates. **j-l,** *CCN1* expression and *GFAP^+^/SLC1A3^+^* astrocytes in healthy human spinal cord grey matter; MS grey matter lesion and adjacent normal appearing grey matter and in human SCI lesion-remote spinal cord grey matter. **m,** Anonymized pathology notes for human tissue used in this study. Scale bar a,b,d,e,f,g: 50 µm, c,j,k,l: 250 µm.

**Extended Data Fig. 10.**
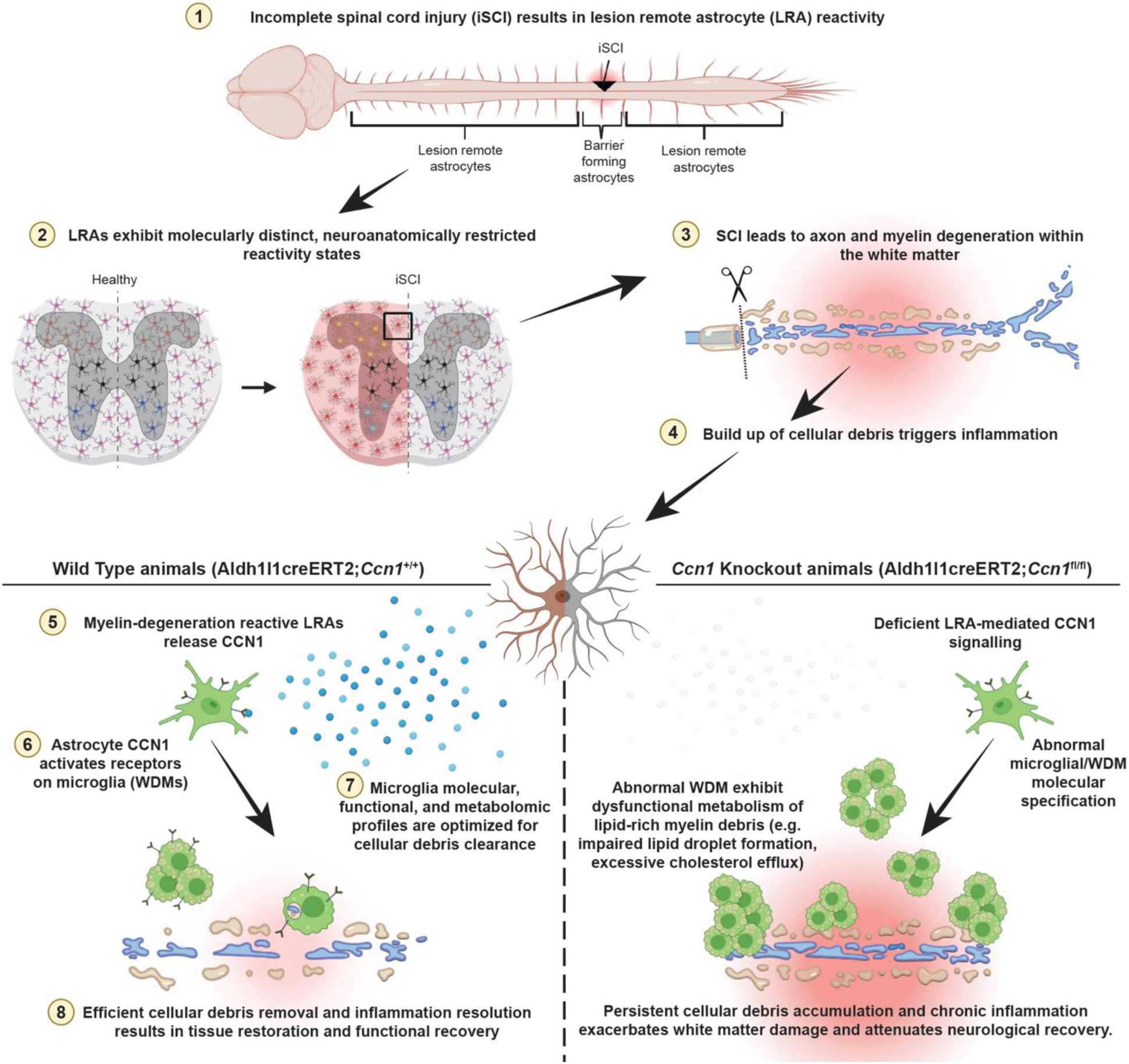
Myelin degeneration-reactive LRAs express CCN1 to regulate local phagocytic microglia and govern white matter repair. Astrocytes in the healthy spinal cord exhibit intraspinal regional molecular heterogeneity. Following SCI, LRA in spared tissue regions exhibit molecularly distinct, neuroanatomically restricted reactivity states that evolve over time. Degeneration of severed axons extends white matter pathology into lesion-remote regions of the injured spinal cord where. In response to local myelin breakdown, reactive white matter LRAs rapidly and persistently upregulate expression of the secreted matricellular signaling protein CCN1. Astrocyte-secreted CCN1 directs the specification and function of white matter degeneration-associated microglia (WDM), which acquire a repair-associated molecular profile and phagocytose myelin and axon debris. In the injured *Ccn1*cKO spinal cord, deficient CCN1 signaling leads to excessive, aberrant activation of WDMs with (i) abnormal molecular specification, (ii) dysfunctional myelin debris processing, and (iii) impaired lipid metabolism, culminating in impaired debris clearance and attenuated functional recovery. Together, these results show that a molecularly and regionally distinct subtype of reactive LRAs plays a critical role in governing white matter inflammation and functionally meaningful neural repair after CNS injury.

